# Visualizing orientation-specific relaxation-diffusion features mapped onto orientation distribution functions estimated via nonparametric Monte Carlo MRI signal inversion

**DOI:** 10.1101/2020.05.23.111963

**Authors:** João P. de Almeida Martins, Chantal M. W. Tax, Alexis Reymbaut, Filip Szczepankiewicz, Derek K. Jones, Daniel Topgaard

## Abstract

Diffusion MRI techniques are widely used to study *in vivo* changes in the human brain connectome. However, to resolve and characterise white matter fibres in heterogeneous diffusion MRI voxels remains a challenging problem typically approached with signal models that rely on prior information and restrictive constraints. We have recently introduced a 5D relaxation-diffusion correlation framework wherein multidimensional diffusion encoding strategies are used to acquire data at multiple echo-times in order to increase the amount of information encoded into the signal and ease the constraints needed for signal inversion. Nonparametric Monte Carlo inversion of the resulting datasets yields 5D relaxation-diffusion distributions where contributions from different sub-voxel tissue environments are separated with minimal assumptions on their microscopic properties. Here, we build on the 5D correlation approach to derive fibre-specific metrics that can be mapped throughout the imaged brain volume. Distribution components ascribed to fibrous tissues are resolved, and subsequently mapped to a dense mesh of overlapping orientation bins in order to define a smooth orientation distribution function (ODF). Moreover, relaxation and diffusion measures are correlated to each independent ODF coordinate, thereby allowing the estimation of orientation-specific relaxation rates and diffusivities. The proposed method is tested on a healthy volunteer, where the estimated ODFs were observed to capture major WM tracts, resolve fibre crossings, and, more importantly, inform on the relaxation and diffusion features along distinct fibre bundles. If combined with fibre-tracking algorithms, the methodology presented in this work may be useful for investigating the microstructural properties along individual white matter pathways.

## 1 INTRODUCTION

For decades, neuroscience has focused not only on unravelling the function of brain areas (Brodmann, 1909; Zilles & Amunts, 2010), but also on the communication between these areas (Sporns, Tononi, & Kötter, 2005). Brain function fundamentally depends on the effective transport of information (Passingham, Stephan, & Kötter, 2002), and inter-individual functional differences may well arise from differences in composition and structure of the physical connections (Alexander-Bloch, Giedd, & Bullmore, 2013). Studying the constituents of the white matter (WM) that form these connections is therefore of paramount importance to understanding brain function in health, disease and development.

The advent of diffusion MRI techniques, which can probe structures at much smaller scales than the imaging resolution by virtue of sensing the random motion of water molecules, has undoubtedly increased the interest in studying WM in the living brain. The possibility to derive quantitative features sensitive to tissue microstructure (Basser & Pierpaoli, 1996; Le Bihan, 1995), and to virtually reconstruct brain connections with fibre-tracking algorithms (Basser, Pajevic, Pierpaoli, Duda, & Aldroubi, 2000; Mori, Crain, Chacko, & Van Zijl, 1999) led to the quick adoption of diffusion MRI in many clinical research applications (Barnea-Goraly et al., 2004; Lebel, Walker, Leemans, Phillips, & Beaulieu, 2008; Lim et al., 1999; Werring, Clark, Barker, Thompson, & Miller, 1999). More recently, tractometry techniques have been developed to tease out WM pathways and characterize their individual tissue microstructure by mapping sets of diffusion-derived parameters along the extracted tracks (Bells et al., 2011; Chamberland et al., 2019; De Santis, Drakesmith, Bells, Assaf, & Jones, 2014; Rheault, Houde, & Descoteaux, 2017; Yeatman, Dougherty, Myall, Wandell, & Feldman, 2012). Fibre-tracking techniques typically rely on the estimation of a fibre Orientation Distribution Function (ODF) per voxel, which is a function on the unit sphere representing the fraction of fibres in each direction (Dell’Acqua & Tournier, 2019; Tournier, 2019). It should be noted that the influence of the fibre ODF is distinct from the orientation distribution of the diffusion signal, and its extraction relies on assessing how tissue microstructure influences the measured MRI signal.

Diffusion MRI studies of WM commonly assume that the voxel-level microstructural features can be adequately represented by a single signal response function (Dell’Acqua & Tournier, 2019; Novikov, Fieremans, Jespersen, & Kiselev, 2019). Under this assumption, the measured signal is written as the convolution between the fibre ODF and a kernel describing the signal response of a set of fibres with a common orientation. The simultaneous unconstrained estimation of the ODF and the microstructural kernel, however, has shown to be notoriously challenging for the diffusion MRI protocols typically used for in vivo research studies (Jelescu, Veraart, Fieremans, & Novikov, 2016). The complexity of this problem is commonly reduced by imposing a set of priors and constraints. Spherical deconvolution of the diffusion MRI signal (Anderson, 2005; Dell’Acqua et al., 2007; Dell’Acqua & Tournier, 2019; Jian & Vemuri, 2007; Tournier, Calamante, & Connelly, 2007; Tournier, Calamante, Gadian, & Connelly, 2004), for example, determines an empirical kernel for the whole brain representing the signal response of a single-fibre population and subsequently solves for the ODF. For heterogeneous voxels containing not only WM but also unknown amounts of grey matter (GM), cerebrospinal fluid (CSF), or pathological tissue, this approach can yield biased ODF estimates.

Multi-tissue spherical deconvolution (Jeurissen, Tournier, Dhollander, Connelly, & Sijbers, 2014) has been proposed to simultaneously resolve sub-voxel tissue fractions and the fibre ODF. While this technique can be used to separate the sub-voxel signal contributions from WM, GM, and CSF, it still assumes a single kernel for all voxels of a given tissue type, which needs to be calibrated *a priori* (Tax, Jeurissen, Vos, Viergever, & Leemans, 2014). Inaccuracies of the calibrated kernels can further bias the estimated fractions and fibre ODFs (Guo et al., 2019; Parker et al., 2013). Alternatively, the voxel-wise kernel can be estimated by first factoring out the ODF through the computation of rotational invariants, and then fitting the data to signal models that set a pre-defined number of microscopic environments with potentially constrained diffusion properties (Kaden, Kelm, Carson, Does, & Alexander, 2016; Novikov et al., 2019; Novikov, Veraart, Jelescu, & Fieremans, 2018). However, different fibre populations within a voxel likely exhibit different microstructural properties (Aboitiz, Scheibel, Fisher, & Zaidel, 1992; De Santis, Assaf, Jeurissen, Jones, & Roebroeck, 2016; Howard et al., 2019; Scherrer et al., 2016), which cannot be reflected with a single voxel-wise kernel. It should furthermore be noted that transverse relaxation time *T*_2_ differences between distinct tissue types are often ignored, which can further bias the quantification of tissue fractions with a single fibre response kernel. The possible existence of a variation of *T*_2_ in anisotropic structures with respect to the orientation of the main magnetic field ***B***_0_ (Lindblom, Wennerström, & Arvidson, 1977), well known in studies of cartilage structure (Henkelman, Stanisz, Kim, & Bronskill, 1994) and more recently reported in in vivo human WM studies (Gil et al., 2016; Knight, Wood, Couthard, & Kauppinen, 2015; McKinnon & Jensen, 2019), would introduce an additional *T*_2_ dispersion and further complicate the quantification of sub-voxel signal fractions. The possible existence of *T*_2_ differences between distinct fibre bundles has motivated the recent development of methods allowing for the measurement of fibre-specific estimates of the transverse relaxation time (de Almeida Martins & Topgaard, 2018; Ning, Gagoski, Szczepankiewicz, Westin, & Rathi, 2020; Schiavi et al., 2019).

Inspired by multidimensional solid-state NMR methodology (Schmidt-Rohr & Spiess, 1994; Topgaard, 2017), we have introduced a framework to quantify the composition of each voxel with joint distributions of effective transverse relaxation rates *R*_2_ = 1/*T*_2_ and apparent diffusion tensors **D** (de Almeida Martins et al., 2020; de Almeida Martins & Topgaard, 2018). Specifically, the inclusion of diffusion MRI data measured with multidimensional diffusion encoding schemes (Topgaard, 2017) and different echo times was observed to alleviate the constraints needed to resolve sub-voxel tissue heterogeneity (de Almeida Martins et al., 2020). Capitalizing on these acquisitions, we quantified sub-voxel compositions using 5D discrete *R*_2_-**D** distributions retrieved from the data using a nonparametric Monte Carlo inversion procedure. However, visualising the retrieved sub-voxel information is challenging because of the high dimensionality of the distributions.

The challenge of visualizing the intricate and comprehensive information within diffusion MRI datasets is an active area of research (Leemans, 2010; Schultz & Vilanova, 2019) and very well established visualization strategies exist to either convey the tensorial properties of a single voxel-averaged **D** (Kindlmann, 2004; Pajevic & Pierpaoli, 2000; Westin et al., 1999) or to visualize a continuous ODF (Peeters, Prckovska, Almsick, Vilanova, & Romeny, 2009; Schultz & Kindlmann, 2010; Tournier et al., 2004; Tuch et al., 2002). However, such techniques are not immediately applicable to the discrete multi-component distributions retrieved with our 5D correlation framework. Previously, we converted the retrieved distributions to sets of statistical parameter maps derived from either the entirety or sub-divisions (‘bins’) of the distribution space (de Almeida Martins et al., 2020; Topgaard, 2019). In ref. (de Almeida Martins et al., 2020), the *R*_2_-**D** space was divided into three bins capturing different ranges of **D** eigenvalues in order to separate the signal contributions from microscopic tissue environments with distinct diffusion properties. Even though bin-resolved maps of signal fractions and means were observed to be useful to map sub-voxel heterogeneity throughout the imaged brain volume, they do not provide information on orientation-resolved properties. In this contribution, we demonstrate how *R*_2_-**D** distributions can be used to derive and visualize fibre-specific relaxation and diffusion metrics. This is done by extending the binning procedure to the space of **D** orientations, and mapping discrete *P*(*R*_2_,**D**) components to a spherical mesh representing a dense set of orientation bins. The attained orientation-resolved information is then conveyed as maps of ODF glyphs that are colour-coded according to the underlying relaxation and diffusion properties; this greatly facilitates the inspection and interpretation of the orientational variation of the 5D *P*(*R*_2_,**D**). The representation of the high-dimensional information as colour-coded ODFs is furthermore compatible with tractography algorithms which hence allows the extension to visualisation of longer-range properties in 3D.

## 2 METHODS

### 2.1 Estimation of 5D relaxation-diffusion distributions

In diffusion MRI, heterogeneous tissues can be described as a collection of microscopic tissue environments wherein water diffusion is modelled by a local apparent diffusion tensor **D**. Within this multi-tensor approach, the diffusion MRI signal is approximated as a weighted sum of the signals from the individual microscopic tissue environments (Jian, Vemuri, Özarslan, Carney, & Mareci, 2007; Novikov et al., 2019; Westin et al., 2016). A similar heterogeneous description has also been used in *R*_2_ studies of intra-voxel brain tissue structure (Does, 2018; MacKay et al., 2006). The transverse relaxation signal of water within tissues is typically expressed as a multi-exponential decay, given by the Laplace transform of a probability distribution of *R*_2_ values (Kroeker & Mark Henkelman, 1986; Whittall & MacKay, 1989; Whittall et al., 1997). Each coordinate of the relaxation probability distribution characterizes the signal fraction of the microscopic environment with the corresponding *R*_2_ rate. Combining the relaxation and diffusion descriptions, the detected signal *S*(*τ*_E_,**b**) can be written as

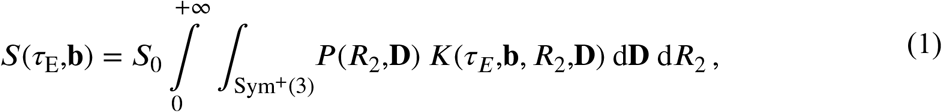

where *P*(*R*_2_,**D**) is the continuous joint probability distribution of *R*_2_ and **D**, *τ*_E_ denotes the echo-time, **b** is the diffusion-encoding tensor, and *S*_0_ is the signal amplitude at vanishing relaxation- and diffusion-weighting, *i.e. S*_0_ = *S*(*τ*_E_ = 0, **b** = 0). The integration of **D** spans over the space Sym^+^(3) of symmetric positive semi-definite 3×3 tensors. The kernel *K*(*τ*_E_,**b**,*R*_2_,**D**) encapsulates the signal decays mapping the distribution onto the detected signal.

Constraining the integral in Eq. (1) to the space of axisymmetric diffusion tensors, each **D** can be parameterized by its axial and radial diffusivities, *D*_∥_ and *D*_⊥_, and by the polar and azimuthal angles, *θ* and *ϕ*, that define its orientation. The *D*_∥_ and *D*_⊥_ eigenvalues can in turn be combined to define measures of isotropic diffusivity *D*_iso_ = (*D*_∥_ + 2*D*_⊥_)/3 and normalized diffusion anisotropy *D*_Δ_ = (*D*_∥_ – *D*_⊥_)/(3*D*_iso_) (Eriksson, Lasič, Nilsson, Westin, & Topgaard, 2015). Using the popular approach of approximating the signal from each microscopic environment as an exponential decay (Dell’Acqua et al., 2007; Does, 2018; Jian et al., 2007; Kaden et al., 2016; MacKay et al., 2006; Novikov et al., 2018; Scherrer et al., 2016; Tuch et al., 2002; Veraart, Novikov, & Fieremans, 2018; Westin et al., 2016), considering only axisymmetric **b**, and adopting the (*D*_iso_,*D*_Δ_,*θ*,*ϕ*) parametrization, Eq. (1) can be expanded as (de Almeida Martins & Topgaard, 2018)

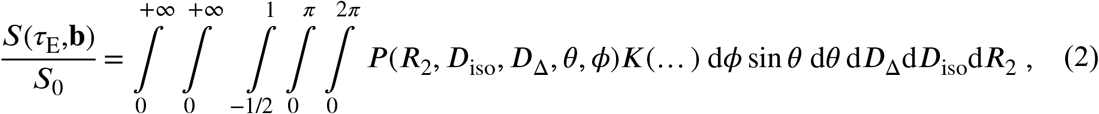

with

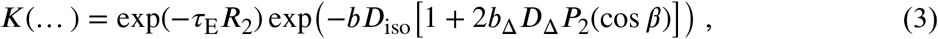

where *b* = Tr(**b**) is recognized as the traditional *b*-value and *b*_Δ_ denotes the normalized anisotropy of the diffusion-encoding tensor (Eriksson et al., 2015). *P*_2_(*x*) = (3*x*^2^-1)/2 is the second Legendre polynomial, and *β* is the shortest angle between the symmetry axes of **D** and **b**. Note that each diffusion orientation (*θ*,*ϕ*) is associated to its own set of microscopic properties (*R*_2_,*D*_iso_,*D*_Δ_) and that no overarching microstructural kernel or universal orientation structure is assumed. This means that Eq. (2) allows for fibre populations with distinct *R*_2_-**D** properties.

For numerical implementation, Eq. (2) is discretized as ***s*** = **K*w***, where ***s*** is the column vector of signal amplitudes measured with *M* combinations of (*τ*_E_,**b**) values, **K** is the inversion kernel matrix, and ***w*** is a vector containing the weights *w_n_* of *N* discrete (*R*_2,*n*_,*D*_∥,*n*_,*D*_⊥,*n*_,*θ_n_*,*ϕ_n_*) configurations. The estimation of ***w*** can then be cast as a constrained linear least-squares problem

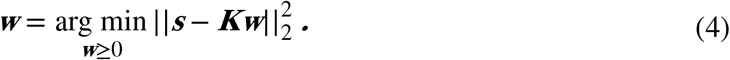

In practice, the argument-minimum operator in Eq. (4) is replaced by a softer condition that searches for a solution within the noise variance. While seemingly straightforward, the problem of finding a solution whose residuals are compatible with the experimental noise is a poorly conditioned one. Indeed, multiple distinct solutions can be found to fit a single noisy dataset. This has motivated the development of several regularization strategies in order to improve the stability of the inverse problem (Daducci et al., 2015; Mitchell, Chandrasekera, & Gladden, 2012; Provencher, 1982; Whittall & MacKay, 1989). A common strategy is to incorporate a regularization term that promotes either a smooth (Benjamini & Basser, 2017; Provencher, 1982; Slator et al., 2019; Venkataramanan, Song, & Hurlimann, 2002) or a sparse (Benjamini & Basser, 2016; Berman, Levi, Parmet, Saunders, & Wiesman, 2013; Urbańczyk, Bernin, Koźmiński, & Kazimierczuk, 2013) ***w*** solution at the expense of a higher residual error.

Monte Carlo algorithms have been used in the porous media field as an alternative to conventional regularized approaches (de Almeida Martins & Topgaard, 2016, 2018; de Kort, van Duynhoven, Hoeben, Janssen, & Van As, 2014; Prange & Song, 2009). These algorithms purposely explore the variability between solutions and estimate an ensemble of distributions consistent with the experimental data. In this work, we use an unconstrained Monte-Carlo algorithm specially designed to handle high-dimensional correlation datasets (de Almeida Martins & Topgaard, 2018; Topgaard, 2019). The algorithm can be broadly divided in two iteration cycles. In the first cycle, the proliferation cycle, a user-defined *N*_in_ number of points is randomly selected from the (log(*R*_2_),log(*D*_∥_),log(*D*_⊥_),cos*θ*,*ϕ*) space, and the corresponding set of weights is found by solving Eq. (4) via a non-negative linear leastsquares algorithm (Lawson & Hanson, 1974); points with non-zero weights are stored and merged with a newly-generated random set. This procedure is repeated for a total of *N*_p_ times, and *N*_p_ random sets of (*R*_2,*n*_,*D*_∥,*n*_,*D*_⊥,*n*_,*θ_n_*,*ϕ_n_*) components are sampled in order to find a configuration yielding sufficiently low residuals. The resulting {(*R*_2,*n*_,*D*_∥,*n*_,*D*_⊥,*n*_,*θ_n_*,*ϕ_n_*)} configuration is stored, duplicated, and its duplicate is then subjected to a small random perturbation. This initiates the second iteration cycle, named mutation cycle, wherein configurations compete with their perturbed counterparts on the basis of lowest sum of squared residuals. The mutation cycle comprises a number of *N*_m_ rounds, following which a possible solution is estimated by selecting the points with the *N* highest weights. In this work we sampled *N*_in_ = 200 points from the (0 < log(*R*_2_ / s^−1^) < 1.5, −11.3 < log(*D*_∥_ / m^2^s^−1^) < −8.3, −11.3 < log(*D*_⊥_ / m^2^s^−1^) < −8.3, 0 < cos*θ* < 1, 0 < *ϕ* < 2π) space, and used *N*_p_ = 20, *N*_m_ =20, and *N* = 20. This inversion was performed voxel-wise and bootstrap with replacement was used in order to estimate per-voxel ensembles of *N*_b_ = 96 solutions, each of which consisting of 20 (*R*_2,*n*_,*D*_∥,*n*_,*D*_⊥,*n*_,*θ_n_*,*ϕ_n_*) components, {(*R*_2,*n*_,*D*_∥,*n*_,*D*_⊥,*n*_,*θ_n_*,*ϕ_n_*)}_1≤*n*≤*N*=20_, and their respective *w_n_* weights.

### 2.2 Resolution of sub-voxel fibre components

Spatially-resolved 5D *R*_2_-**D** distributions were estimated using the procedure described in the previous section. As the main brain components – white matter (WM), grey matter (GM), and cerebrospinal fluid (CSF) – are characterized by clearly distinct diffusion properties, we expect most (*R*_2,*n*_,*D*_∥,*n*_,*D*_⊥,*n*_,*θ_n_*,*ϕ_n_*) components to agglomerate within three distant regions of the diffusion space (Pierpaoli, Jezzard, Basser, Barnett, & Di Chiro, 1996).

The idea that most *P*(*R*_2_,**D**) components will fall within three coarse regions has inspired the division of the *R*_2_-**D** space into three smaller subsets (‘bins’) based on the diffusion properties of WM, GM, and CSF (de Almeida Martins et al., 2020). We then defined three bins named ‘thin’ (0.6 < log(*D*_∥_/*D*_⊥_) < 3.5, −10 < log(*D*_iso_/m^2^s^−1^) < −8.7, −0.5 < log(*R*_2_/s^−1^) < 2), ‘thick’ (–3.5 < log(*D*_∥_/*D*_⊥_) < 0.6, −10 < log(*D*_iso_/m^2^s^−1^) < −8.7, −0.5 < log(*R*_2_/s^−1^) < 2), and ‘big’ (–3.5 < log(*D*_∥_/*D*_⊥_) < 3.5, −8.7 < log(*D*_iso_/m^2^s^−1^) < −8, −0.5 < log(*R*_2_/s^−1^) < 2). The names of the different bins highlight the geometry of the **D** captured by each one of them. For each bootstrap realization *n*_b_ (1 ≤ *n*_b_ ≤ *N*_b_), signal contributions from anisotropic tissues are resolved by selecting the set of *P*(*R*_2_,**D**) components that fall within the ‘thin’ bin:

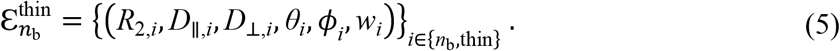

The {(*R*_2,*i*_,*D*_∥,*i*_,*D*_⊥,*i*_,*θ_i_*,*ϕ_i_*,)}_*i*∈ {*n*_b_,thin}_ configurations and {*w_i_*}_*i*∈ {*n*_b_,thin}_ weights of 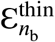 are interpreted as representing the *R*_2_-**D** properties and signal fractions, respectively, of a discrete set of sub-voxel fibre populations. The binning and anisotropic selection processes are illustrated in panels A and B of **Figure 1**.

**Figure 1.**
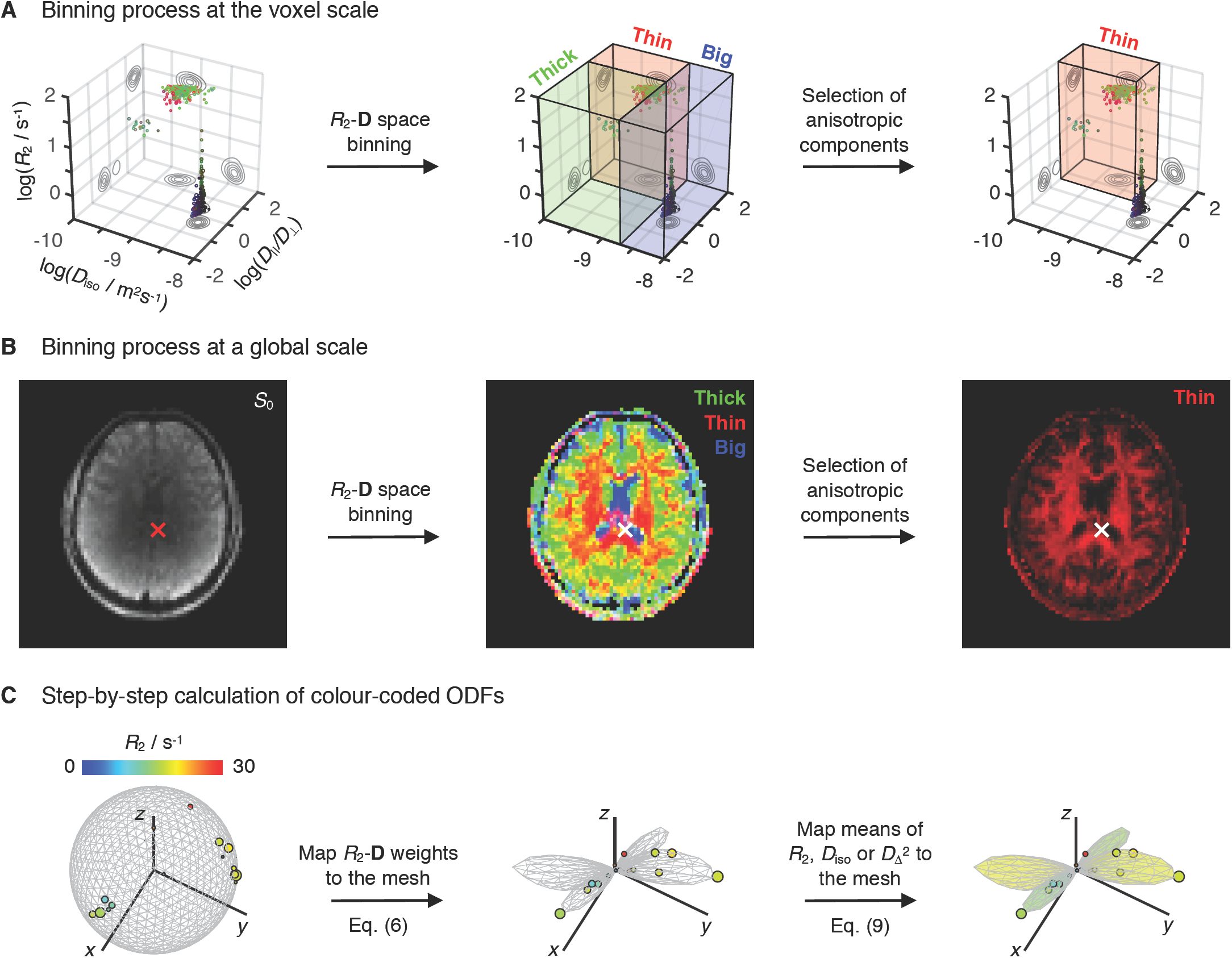
Resolution of sub-voxel fibre-like components and subsequent estimation of the associated colour-coded Orientation Distributions Functions (ODFs). (A) *R*_2_-**D** distribution obtained for a voxel containing both CSF and two crossing WM populations. The 5D *P*(*R*_2_,**D**) is reported as a 3D logarithmic scatter plot of *R*_2_, isotropic diffusivities *D*_iso_, and axial-radial diffusivity ratios *D*_∥_/*D*_⊥_, with circle area proportional to the weight of the corresponding *R*_2_-**D** component, *w*. Colour coding is defined as: [R,G,B] = [cos*ϕ* sin*θ*, sin*ϕ* sin*θ*, cos*θ*] *×* |*D*_∥_-*D*_⊥_|/max(*D*_∥_, *D*_⊥_), where (*θ*,*ϕ*) gives the orientation of each axisymmetric **D**. The *R*_2_-**D** space is divided into three coarse bins named ‘big’ (blue volume), ‘thin’ (red volume), and ‘thick’ (green volume). Components falling in the ‘thin’ bin are singled-out and interpreted as fibres. (B) Spatial distribution of per-bin signal contributions. The middle map shows the fractional populations in the ‘big’ (blue), ‘thin’ (red), and ‘thick’ (green) bins as a colour-coded composite image. The rightmost map focuses on the signal contributions from components within the ‘thin’ subset, *f*_thin_, the complement of which, (1 – *f*_thin_), gives the signal fraction from all components not used for ODF calculation. The crosses locate the voxel whose distribution is shown in panel (A). (C) Scheme for calculating colour-coded ODFs. The *R*_2_-coloured circles denote the ‘thin’ components from a bootstrap solution of the voxel signalled in panel (B). Circle area is proportional to *w*, while the [*x*,*y*,*z*] circle coordinates are defined as either [cos*ϕ* sin*θ*, sin*ϕ* sin*θ*, cos*θ*] (left) or [cos*ϕ* sin*θ*, sin*ϕ* sin*θ*, cos*θ*] · *w* (middle and right). In the left plot, the discrete *R*_2_-**D** components are displayed on a unit sphere represented by a 1000-point (*θ*,*ϕ*) mesh. The weights of the *P*(*R*_2_,**D**) components are first mapped to the mesh through Eq. (6), creating an ODF glyph whose radii inform on the *R*_2_-**D** probability density along a given (*θ*,*ϕ*) direction (middle). Following the ODF estimation, Eq. (9) is used to assign mean values of *R*_2_, *D*_iso_, or *D*_Δ_ to each mesh point and define the colour the ODF glyph (right).

### 2.3 ODF estimation

The colour-coded 3D scatter plots of *R*_2_, *D*_iso_, and *D*_∥_/*D*_⊥_ displayed in **Figure 1** allow the visualization of the full set of properties of the voxel-wise 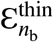 components. Despite its usefulness, the scatter plot representation concentrates points corresponding to anisotropic components within a small region of the (*R*_2_, *D*_iso_, *D*_∥_/*D*_⊥_) space, which in turn makes it difficult to evaluate their orientation properties in detail. For example, while **Figure 1A** clearly informs on the existence of two fibre populations oriented along two different directions (red and green points), it does not provide unambiguous information about the relative signal contributions of the two populations. To better inspect the orientational information of the underlying *P*(*R*_2_,**D**), it is helpful to convert the discrete set of fibre orientations to a continuous object informing on the *R*_2_-**D** probability density in each direction, which can then be visualized as a single glyph with an intuitive geometrical interpretation. To achieve that, we used a 1000-point triangle mesh on the unit sphere, created via an electrostatic repulsion scheme (Bak & Nielsen, 1997; Jones, Horsfield, & Simmons, 1999), to create 1000 orientation bins; the mesh vertices define the centres of (*θ*,*ϕ*) bins without clearly defined boundaries. The median angular distance between nearest-neighbouring mesh points (or bin centres) is approximately 7°. Afterwards, a smoothing kernel was used to map the weights of 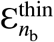 onto the dense set of (*θ,ϕ*) bins. The role of the smoothing kernel is to weight the influence of each bin according to the angular distance between its centre and a given 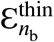 component, and to distribute the contributions of each discrete {*w_i_*}_*i*∈{*n*_b_,thin}_ throughout various bins in order to define a smooth Orientation Distribution Function (ODF) *P*_*n*_b__(*θ*,*ϕ*) that can be straightforwardly visualized as a polar plot (Leemans, 2010; Schultz & Vilanova, 2019).

In this work, the orientation binning and consequent estimation of *P*_*n*_b__(*θ*,*ϕ*) was performed through a convolution with a Watson distribution kernel (Mardia & Jupp, 2009; Watson, 1965):

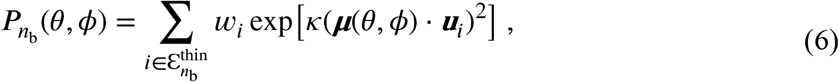

where ***u**_i_* is the unit vector describing the orientation of the *i*-th discrete component and ***μ***(*θ*,*ϕ*) is the unit mean direction vector of bin centre (*θ*,*ϕ*). The variable *κ* denotes a concentration parameter that regulates the amount of dispersion about ***μ***(*θ*,*ϕ*). Further insight onto the nature of the Watson distribution kernel and the role of parameter *κ* is attained by considering the small-angle approximation of the former

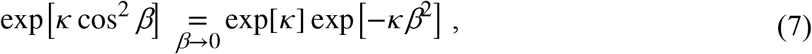

where *β* is the shortest angle between unit vectors ***u*** and ***μ***. Within this approximation, the Watson kernel is rewritten as a familiar Gaussian smoothing kernel whose standard deviation, *σ*, defines an angular spreading that is directly related to the concentration parameter *κ*, *σ* = (2*κ*)^−1/2^. The relationship between *κ* and *σ* enables us to easily set a dispersion factor in relation to the angular distance of the various mesh points. To cover the minimum angular distance between mesh points, we set *σ* to be 50% higher than the median angular distance between nearest-neighbouring mesh points, *i.e*. *σ* = 10.5° or, equivalently, *κ* = 14.9. The rationale behind our choice of *κ* is elaborated upon in section 3.1.

Separate ODFs were calculated for each of the *N*_b_ = 96 bootstrapped *P*(*R*_2_,**D**) solutions; the final *P*(*θ*,*ϕ*) was then estimated as the median of the *N*_b_ independent ODFs:

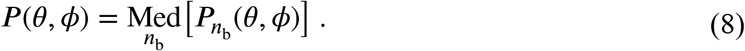

As the orientation of each fibre solution is correlated to a given set of {*R*_2,*n*_,*D*_∥,*n*_,*D*_⊥,*n*_} coordinates, we can assign any statistical descriptor of the (*R*_2_,*D*_iso_,*D*_Δ_) space to the various coordinates of *P*(*θ*,*ϕ*). In line with previous works (de Almeida Martins et al., 2020; Topgaard, 2019) where *R*_2_-**D** were converted into bin-resolved mean maps of *R*_2_, *D*_iso_, and *D*_Δ_^2^, we map mean values of *R*_2_, *D*_iso_, and *D*_Δ_^2^ into the ODF mesh. The mean value of *X* = *T*_2_, *R*_2_, *D*_iso_, or *D*_Δ_^2^ per mesh orientation for each bootstrap *n*_b_, 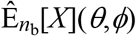, is calculated as

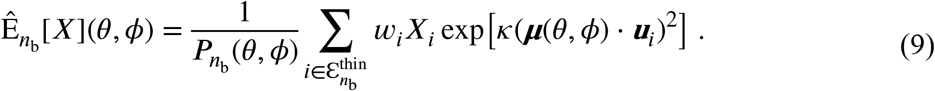

By mapping different descriptors of the *R*_2_-**D** distributions to specific ODF coordinates, we can visualize orientation-specific information on tissue composition and structure. As before, the final voxelwise 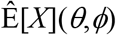 is estimated as the median of the individual per-bootstrap 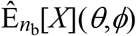 values:

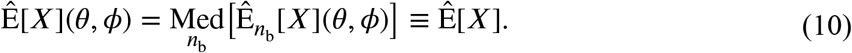

For compactness, we omit the explicit (*θ*,*ϕ*) dependence from the orientation-resolved means and simply denote them as 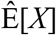. The 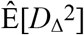 metric provides orientation-resolved information on the underlying mean-diffusivity. The 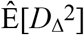 metric is the orientation-resolved counterpart of the mean *D*_Δ_^2^ descriptor (Topgaard, 2019), which is in turn similar to previously introduced anisotropy measures such as the microscopic anisotropy index (MA) (Lawrenz, Koch, & Finsterbusch, 2010), the fractional eccentricity (FE) (Jespersen, Lundell, Sønderby, & Dyrby, 2013), and the microscopic fractional anisotropy (μFA) (Lasič, Szczepankiewicz, Eriksson, Nilsson, & Topgaard, 2014).

The mapping of *P*(*R*_2_,**D**) components to a dense mesh as described by equations (6)–(10) is a key result from this contribution, and provides the basis for extracting and visualizing orientation-resolved information from nonparametric *R*_2_-**D** distributions. **Figure 1C** illustrates how both 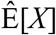 and the associated orientation distribution can be conveniently represented by colour-coded ODF glyphs; the shape of the glyph reflects the *P*(*θ*,*ϕ*) distribution, while the colour informs on the 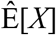 values at the various (*θ*,*ϕ*) points. Functions used to compute the colour-coded ODFs have been incorporated in the multidimensional diffusion MRI toolbox (Nilsson et al., 2018): https://github.com/JoaoPdAMartins/md-dmri. In this work, maps of ODF glyphs were computed using those same functions and rendered with POV-Ray (http://www.povray.org/).

The local maxima of the computed ODFs can be used to estimate “peaks” providing information on the main orientations of the sub-voxel diffusion pattern. The peaks provide a rough and easily accessible measure of the orientation and *R*_2_-**D** properties of different ODF lobes and can consequently be used to assess the properties of the various fibre populations. Here, up to four peaks per voxel were determined by assessing the mesh points (*θ*,*ϕ*) for which *P*(*θ*,*ϕ*) is a local maximum and *P*(*θ*,*ϕ*)/max_(θ,ϕ)_[*P*(*θ*,*ϕ*)] ≥ 0.1. The *R*_2_-**D** properties of the ODF peaks were estimated by calculating 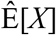 (see Eq. (9)) for each peak orientation.

### 2.4 *In vivo* data acquisition

Multidimensional relaxation-diffusion MRI data were acquired using a prototype spin-echo diffusion weighted sequence with an echo-planar imaging (EPI) readout, customized for diffusion encoding with user-defined gradient waveforms (Lasič et al., 2014; Szczepankiewicz, Sjolund, Stahlberg, Latt, & Nilsson, 2019). Images were recorded with the following parameters: TR = 4 s, FOV = 234×234×60 mm^3^, voxel-size=3×3×3 mm^3^, partial-Fourier = 6/8, and a parallel-imaging (GRAPPA) factor of 2. Diffusion encoding was performed with a set of five gradient waveforms yielding *b*-tensors with four distinct “shapes” (*b*_Δ_ = −0.5, 0.0, 0.5, and 1) (Eriksson et al., 2015). The various waveforms were used to probe *b*-tensors of varying magnitude *b*, anisotropy *b*_Δ_, and orientation (*θ*,*ϕ*) at different echo-times *τ*_E_; this procedure yields 5D relaxation-diffusion correlated datasets whose dimensions match those of the sought-for distributions. Readers interested in the sequence used in this work are directed to a public repository: https://github.com/filip-szczepankiewicz/fwf_seq_resources.

**Figure 2** displays the waveforms used in this work and **Table 1** summarizes the (*τ*_E_,**b**) acquisition protocol. Besides the (*τ*_E_,**b**) points detailed in **Table 1**, we additionally acquired *b* = 0 images with reversed phase-encoding blips at *τ*_E_ = 60, 80, 110, and 150 ms in order to correct for susceptibility-induced distortions (Andersson, Skare, & Ashburner, 2003). The acquisition scheme comprised a total of 686 (*τ*_E_,**b**) acquired over an imaging session of ~45 mins. The asymmetric waveforms giving *b*_Δ_ = −0.5, 0.0, and 0.5 were calculated with a MATLAB package (https://github.com/jsjol/NOW) that optimizes for maximum *b* (Sjölund et al., 2015) and minimizes the effects of concomitant magnetic field gradients (Szczepankiewicz, Westin, & Nilsson, 2019). Linearly encoded (*b*_Δ_ = 1) data was acquired with two different gradient waveforms: a non-monopolar gradient waveform and a standard Stejskal-Tanner waveform (Stejskal & Tanner, 1965). The asymmetric gradient pulses from the non-monopolar waveform were manually designed with the aim of minimizing the differences between the spectral profile of that *b*_Δ_ = 1 waveform and the spectral profile of the *b*_Δ_ ≠ 1 waveforms (Lundell et al., 2019). The Stejskal-Tanner design was used to probe a shorter *τ*_E_ and higher *b*-values (*b* = 4·10^9^ m^−2^s) than those achievable with the non-monopolar *b*_Δ_ = 1 waveform. While the measured apparent diffusivities are known to be related to the frequency spectra of the gradient waveforms (Callaghan & Stepišnik, 1996; Lundell et al., 2019; Stepišnik, 1993), such a relationship is likely to have a negligible effect on healthy human brain data acquired with the limited range of frequency contents probed in this work (Szczepankiewicz, Lasič, et al., 2019) and no biases are expected to originate from the spectral differences of the *b*_Δ_ = 1 waveforms.

**Figure 2.**
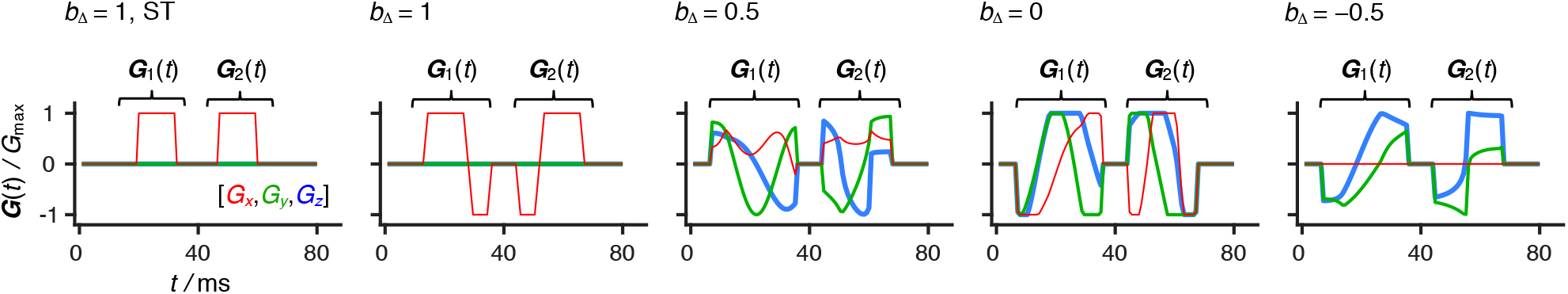
Set of gradient waveforms used in this study. The ST acronym identifies a standard Stejskal-Tanner waveform whose spectral profile (Callaghan & Stepišnik, 1996; Lundell et al., 2019) is distinct from those of the non-monopolar waveforms. The waveforms yielding *b*_Δ_ = –0.5, 0, and 0.5 were optimized according to the numerical procedure described in refs. (Sjölund et al., 2015) and (Szczepankiewicz, Westin, et al., 2019). The displayed waveforms were inserted within a spin-echo sequence with an EPI readout. To clarify the locations of the spin-echo radio-frequency pulses and the EPI block, we divide each waveform in two components: ***G***_1_(*t*) and ***G***_2_(*t*). The 90° pulse is executed at *t* = 0, before the ***G***_1_(*t*) component, while the 180° pulse is applied after a time *t* = *τ*_E_/2 and is bracketed by the ***G***_1_(*t*) and ***G***_2_(*t*) components. Signal readout starts shortly after the conclusion of ***G***_2_(*t*), with the EPI block being centred at the echo-time *τ*_E_. Relaxation-weighting is enforced by varying *τ*_E_ while keeping constant the location of ***G***_1_(*t*) and ***G***_2_(*t*) relative to the 180° pulse.

**Table 1.**
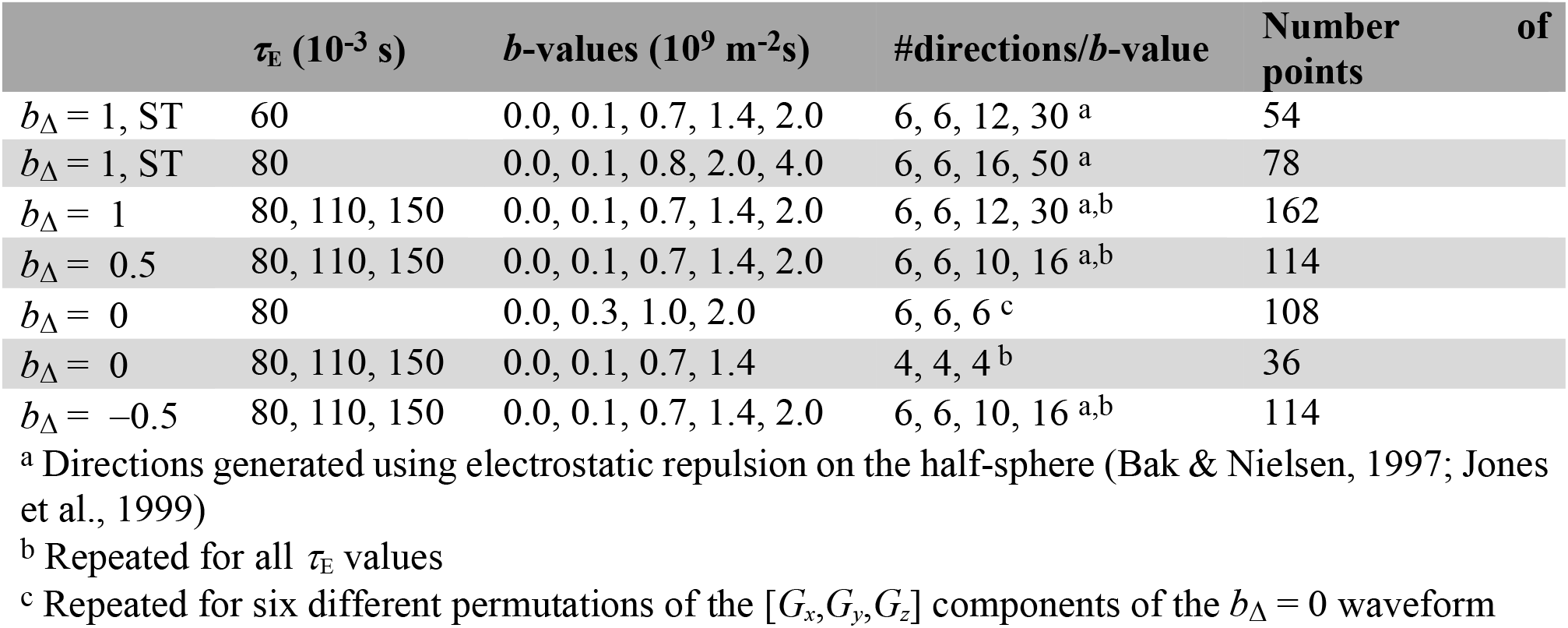
5D relaxation-diffusion correlation protocol used in this work.

The protocol described above was implemented on a 3T Siemens MAGNETOM Prisma scanner (Siemens Healthcare, Erlangen, Germany) and used to scan a healthy adult volunteer. This study was approved by the Cardiff University School of Psychology ethics committee, and informed written consent was obtained prior to scanning.

### 2.5 Post processing

The entire dataset was divided in *τ*_E_-specific data subsets, which were denoised using random matrix theory (Veraart et al., 2016), and corrected for Gibbs ringing artefacts using the method described in (Kellner, Dhital, Kiselev, & Reisert, 2016). Signal drift correction was subsequently performed as detailed in ref. (Vos et al., 2017). The acquired data were further corrected for subject motion and eddy-current artefacts using ElastiX (Klein, Staring, Murphy, Viergever, & Pluim, 2009) with extrapolated references (Nilsson, Szczepankiewicz, van Westen, & Hansson, 2015) as implemented in the multidimensional diffusion MRI toolbox (Nilsson et al., 2018); this procedure was performed with the default settings of the toolbox to the entire (*τ*_E_,**b**) dataset. Susceptibility-induced geometrical distortions were corrected using the TOPUP tool in the FMRIB software library (FSL) (Smith et al., 2004), with the same settings being applied to the entire (*τ*_E_,**b**) dataset.

### 2.6 *In silico* datasets

The angular resolution of our acquisition and analysis protocols was investigated using *in silico* data. We simulated a two-component system designed to mimic two fibres with similar diffusion features but distinct *R*_2_ rates:

- Component 1: *T*_2_ = 1/*R*_2_ = 60 ms, *D*_iso_ = 0.75·10^−9^ m^2^s^−1^, *D*_Δ_ = 0.9, *w* = 0.5;
- Component 2: *T*_2_ = 1/*R*_2_ = 100 ms, *D*_iso_ = 0.75·10^−9^ m^2^s^−1^, *D*_Δ_ = 0.9, *w* = 0.5.

The orientation of component 1 was kept constant (*θ* = 0, *ϕ* = 0), while that of component 2 was varied in order to define four distinct crossing angles: (*θ* = 25°, *ϕ* = 0), (*θ* = 30°, *ϕ* = 0), (*θ* = 35°, *ϕ* = 0), and (*θ* = 40°, *ϕ* = 0).

The ground-truth signal data for the four fibre-crossing systems were generated using the (*τ*_E_,**b**) acquisition scheme indicated in Table 1 and computed from Equation (2). Gaussian distributed noise with an amplitude of 1/SNR was added to the ground-truth signals in order to simulate the effects of experimental noise. The experimental SNR, computed as the mean-to-standard-deviation-ratio of *b*_Δ_ = 0 data acquired at *b* = 0.3·10^9^ m^−2^s and *τ*_E_ = 80 ms (Szczepankiewicz, Sjolund, et al., 2019), was estimated to SNR = 72 ± 28 for WM regions. Consequently, we defined SNR = 70 for the *in silico* calculations, a value that is compatible with the SNR of the *in vivo* data. In line with a recent *in silico* study of the performance of the Monte Carlo algorithm in inverting multidimensional diffusion data (Reymbaut, Mezzani, de Almeida Martins, & Topgaard, 2020), we drew 100 independent noise configurations and computed 100 different signal realizations for each of the four fibre-crossing systems. The various signal realizations were inverted using the Monte-Carlo algorithm described in section 2.1, and the resulting solution ensembles were subsequently compared against the corresponding ground-truth systems.

## 3 RESULTS & DISCUSSION

### 3.1 Defining the dispersion factor of the Watson kernel

As mentioned in section 2.3, the use of a Watson kernel introduces an artificial angular dispersion to the inverted 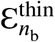 components. The amount of angular dispersion is regulated by the user-defined parameter *κ*, and is introduced to assure that the discrete {*w_i_*}_*i*∈{*n*_b_,thin}_ weights do not fade between the intervals of the ***μ***(*θ*,*φ*) mesh. To understand the smoothing effects of the Watson kernel over a discrete mesh, it is instructive to consider the decay of the Watson function over a given angular distance Δ*β*: *v* = (exp[*κ*cos^2^ Δ*β*] – 1)/ (exp[*κ*] – 1). Considering a 1000-point mesh and *κ* = 14.9, the values used for the *in vivo* data analysis, the maximum distance between an arbitrary {*θ_i_*,*ϕ_i_*,}_*i*∈{*n*_b_,thin}_ configuration and the nearest mesh point is ~3.5°, a value for which the Watson kernel retains *v* = 0.95 of its maximal influence. The minimal decay of the Watson kernel over Δ*β* = 3.5° ensures that the set of 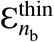 discrete components is indeed mapped into the mesh. From Equation (7) it is additionally obvious that the choice of *κ* is a trade-off between a sufficiently smooth ODF representation and the angular resolution of the ODF in disentangling different peaks. The question then arises whether setting *κ* = 14.9 may over-smooth the orientational information within the *R*_2_-**D** distributions. For instance, with *κ* = 14.9 the Watson kernel will retain more than 50% (*v* = 0.64) of its maximum value over a distance of Δ*β* = 10°, meaning that 20° crossings cannot be resolved with our settings. To assess if the amount of *κ*-generated dispersion is sufficiently low not to misrepresent the orientational information of the *R*_2_-**D** distributions, we investigated *in silico* the angular resolution of the Monte Carlo analysis.

The angular resolution of our framework was assessed by inverting *in silico* data from two anisotropic components crossing at various angles (see section 2.6 for further details). The (*R*_2_,*θ*) projections of the attained *R*_2_-**D** distributions are displayed in **Figure 3A** and inform that, at the SNR of the *in vivo* data, crossings of 30° or higher can be directly resolved in the Monte Carlo *P*(*R*_2_,**D**). Setting equal *R*_2_ (*T*_2_ = 1/*R*_2_ = 60 ms) properties for both anisotropic components or adding a third isotropic component (*T*_2_ = *1/R*_2_ = 500 ms, *D*_iso_ = 2·10^−9^ m^2^s^−1^, *D*_Δ_ = 0) with a total signal fraction of up to 0.2 did not affect the angular resolution of the 30° crossing, but lead to an underestimation of the signal fraction from the *θ* = 30° fibre population. The accurate resolution of the 35° and 40° systems was unaffected by changes in component *R*_2_ or the introduction of the fast-diffusing component. The *in silico* results then suggest that the maximum achievable angular resolution of our experimental protocol is between 30° and 35°, and a conservative approach is to set *κ* so that 35° crossings are not over-smoothed and obscured. Computing ODFs for the *in silico* distributions confirms that setting *κ* = 14.9 is indeed sufficient to resolve a 35° crossing (see **Figure 3B**). While there is room to increase *κ* without risking 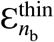 disappearing though the holes of the mesh, we observe that a significantly sharper Watson kernel leads to narrow ODF lobes that do not accurately portray the angular dispersion of the underlying *R*_2_-**D** distributions (compare panels **A** and **B** of **Figure 3**). Moreover, significantly higher *κ* values were tested in the in vivo dataset and observed to lead to non-smooth ‘spiky’ ODFs in voxels containing orientationally dispersed fibres.

**Figure 3.**
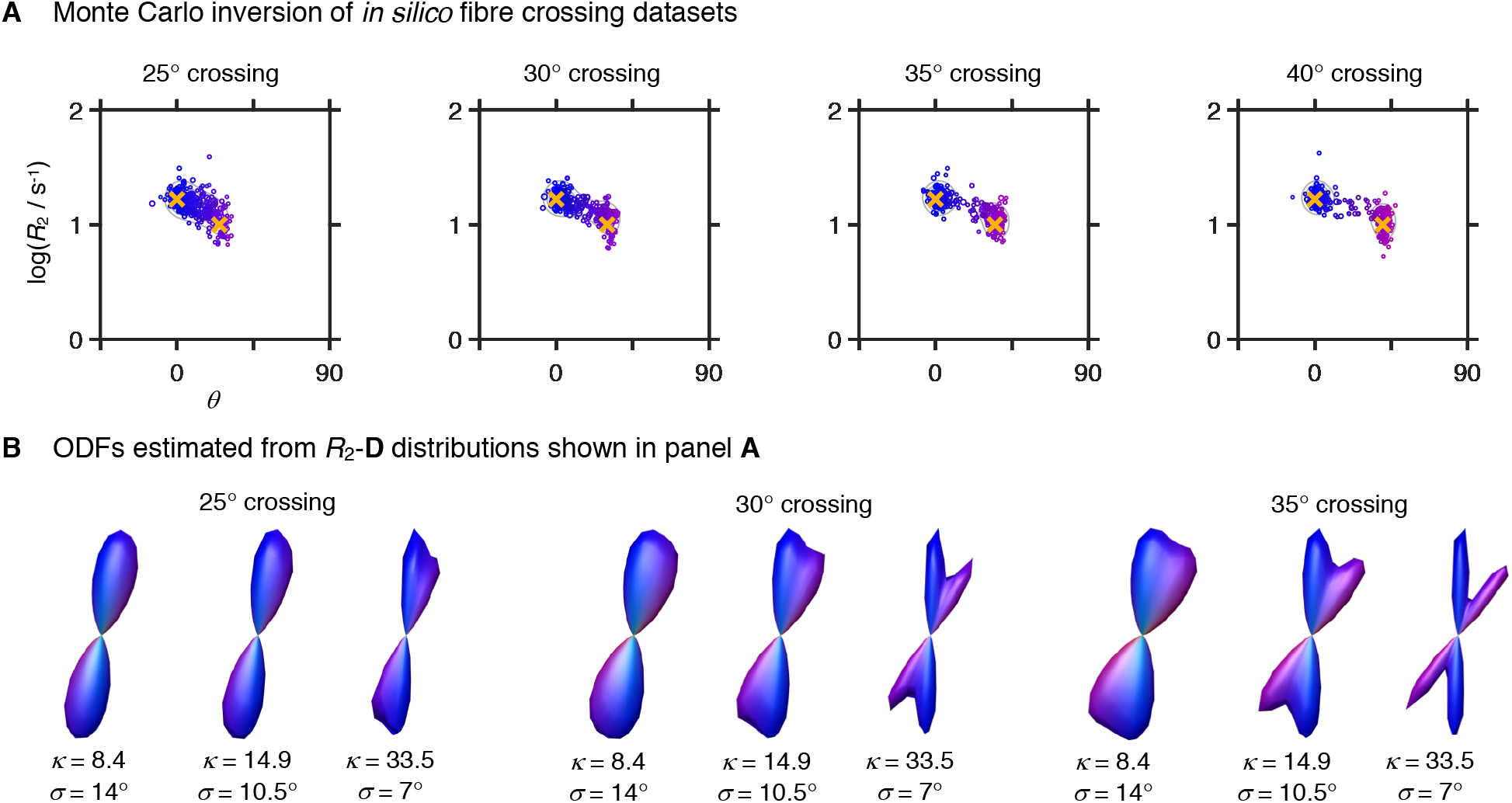
*R*_2_-**D** distributions and Orientation Distribution Functions (ODF) retrieved for *in silico* fibre-crossing datasets. (A) 5D *P*(*R*_2_,**D**) distributions displayed as 2D scatter plots of log(*R*_2_) and *θ*, the polar angle defining **D** orientation. Circle area is proportional to the weight of the corresponding component and colouring is defined as [R,G,B] = [cos*ϕ* sin*θ*, sin*ϕ* sin*θ*, cos*θ*] · |*D*_∥_–*D*_⊥_|/max(*D*_∥_, *D*_⊥_), where *D*_∥_ and *D*_⊥_ denote the axial and radial diffusivities, respectively, and *ϕ* is the azimuth angle of **D**. The yellow crosses identify the ground-truth values. (B) ODF glyphs estimated from the distributions in panel (A), using Watson kernel with different orientation dispersion factors (see Eq. (6) for further details). The ODF colouring follows a conventional directional scheme: [R,G,B] = [*μ_xx_*,*μ_yy_*,*μ*_zz_], where *μ_ii_* are the elements of the unit vector ***μ***(*θ*,*ϕ*) defining the orientation of mesh-point (*θ*,*ϕ*).

### 3.2 *In vivo* fibre orientations

Previous work from our group (de Almeida Martins et al., 2020) has shown that pure component voxels containing either WM, GM, or CSF give rise to clearly distinctive *R*_2_-**D** distributions that accurately capture the main microscopic features of the various tissues – CSF: high isotropic diffusivity *D*_iso_, low normalized diffusion anisotropy *D*_Δ_, low *R*_2_; WM: low *D*_iso_, high *D*_Δ_, high *R*_2_; GM: low *D*_iso_, low *D*_Δ_, high *R*_2_. Voxels comprising mixtures of GM, WM, and GM are in turn characterized by multimodal distributions that exhibit a linear combination of properties of the distributions from the individual components. **Figure 1A** displays the distribution obtained from a voxel containing both CSF and contributions from two WM tracts: the *corpus callosum* (CC) and the *fornix*. Three distinct tissue environments can be clearly discerned in the displayed distribution: an isotropic fast diffusing component attributed to CSF and two anisotropic slow diffusing components with different orientations corresponding to the WM tracts. By ascribing distribution points to one of the three bins discussed in the Methods section we were able to separate and quantify the signal contributions from distinct brain tissues. Indeed, as shown in **Figure 1B**, the signal fractions from the various bins follow the expected spatial distributions of WM, GM, and CSF.

**Figure 4** displays the ODFs computed from the components that fall within the ‘thin’ bin. The ODFs are displayed as directionally-coloured glyphs, superimposed on the sum of the signal fractions from the ‘big’ and ‘thick’ populations. Overall, the reconstructed ODFs are consistent with the expected WM arrangement of the healthy human brain. Major WM tracts such as the corticospinal tract (CST), the CC, and the superior longitudinal fasciculus (SLF) are easily located in the displayed figure (see arrows in **Figure 4**), and multiple crossings can also be discerned. The zoomed panels show that the proposed method can capture the crossings in the ventral SLF – anterior-posterior fibres with leftright fibres – and the crossings between the CST and the CC – superior-inferior fibres with left-right fibres. The dotted boxes show that three-fibre crossings present in the *centrum semiovale* are well captured by this technique, meaning that more than two fibre populations can be resolved.

**Figure 4.**
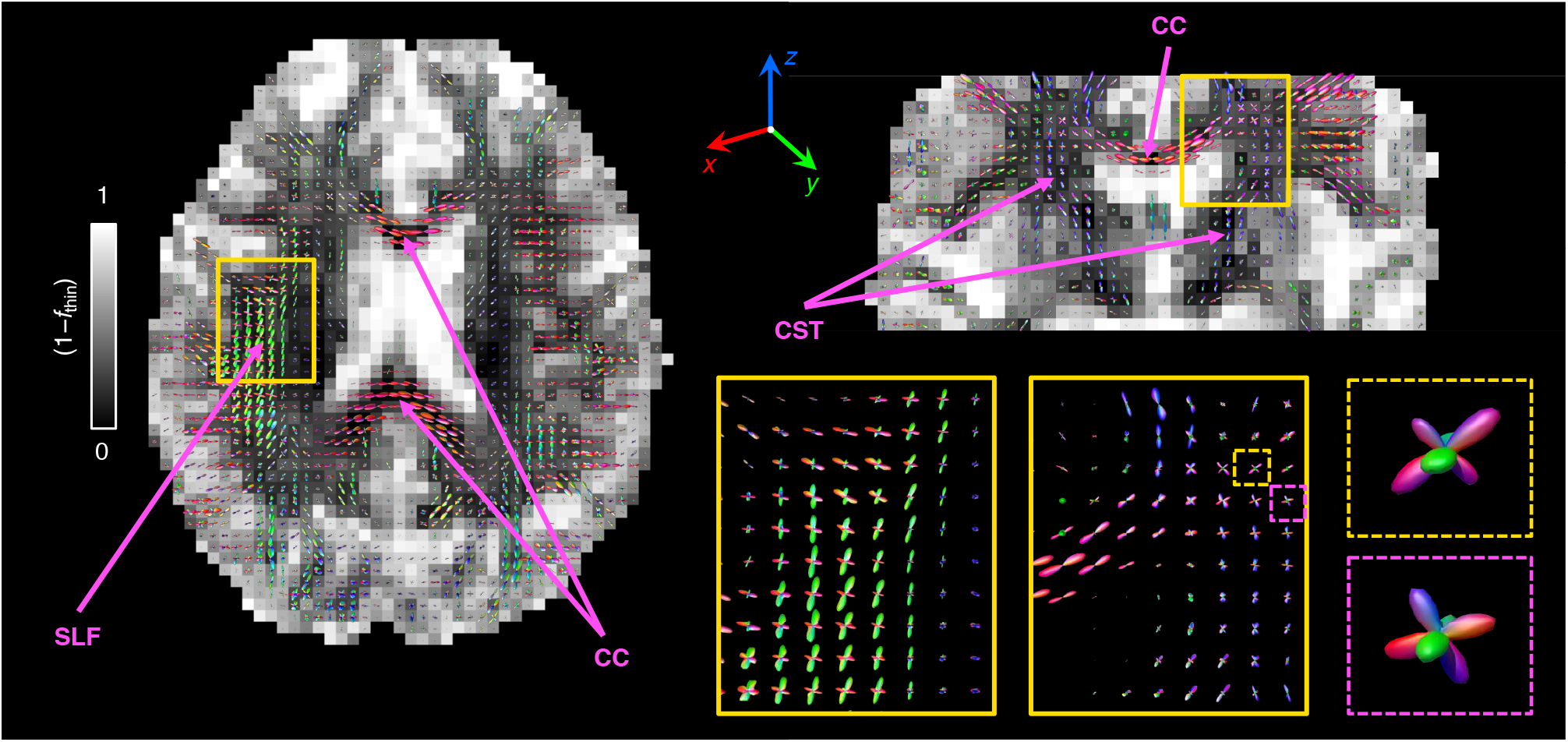
Per-voxel Orientation Distribution Functions (ODF), *P*(*θ*,*ϕ*), estimated from *R*_2_-**D** distribution components ascribed to the ‘thin’ bin defined in **Figure 1**. The voxel-wise *P*(*θ*,*ϕ*) were computed by using Eq. (6) to map the weights of the bin-resolved discrete *P*(*R*_2_,**D**) components into a 1000-point spherical mesh. Here, each ODF is represented as a 3D polar plot with a local radius given by *P*(*θ*,*ϕ*) and colour-coded according to [R,G,B] = [*μ_xx_*,*μ_yy_*,*μ*_zz_], where *μ_ii_* are the elements of the unit vector ***μ***(*θ*,*ϕ*) (see Eq. (6)) for further details). In the left and top-right panels, the sets of ODF glyphs are superimposed on a grey-scaled map that shows the signal contributions from non-fibre-like components (1–*f*_thin_), *i.e*., signal fractions from the ‘big’ and ‘thick’ populations. The zoom-ins in the lower-right panel offer a more detailed look into selected fibre crossing regions (continuous line boxes) and three-fibre crossing voxels (dashed line boxes) found in the *centrum semiovale*. The various arrows identify fibre tracts mentioned in the main text.

Voxels at the WM-CSF and WM-GM interfaces exhibit small-amplitude ODFs, consistent with lower signal fractions of fibrous tissue. The low amplitude of the ODF lobes found in those regions does not seem to bias their orientation; for example, CC voxels near the ventricles yield low amplitude lobes whose orientations follow the expected trend (fibres running left-right). These observations indicate that the estimated ODFs are robust to partial volume effects with CSF and that the proposed method can indeed resolve fibre orientations in heterogeneous voxels. *In silico* calculations show that an accurate ODF can be estimated as long as the contribution from CSF accounts for less than 75% of the total voxel-signal. Low-amplitude ODF lobes can also be found throughout cortical GM regions. These ODFs might be explained by the presence of anisotropic tissue components in cortical GM (Assaf, 2019), or interpreted as originating from low-amplitude WM partial volume effects caused by the large voxel-size used in this study. A more in-depth study than the one presented in this contribution is necessary in order to unambiguously discriminate between these two factors.

### 3.3 *In vivo* orientation-resolved *R*_2_-D metrics

The relaxation and diffusion features from different fibres can be investigated by using Eq. (9) to map *R*_2_-**D** metrics onto the ODF mesh and define orientation resolved means, 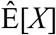. The estimated 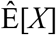 values are then visualized as colour-coded ODF glyphs such as the ones displayed in **Figure 5**, which inform on the correlations between **D** orientation and *R*_2_, *D*_iso_, or *D*_Δ_^2^. The displayed ODF maps capture the expected diffusion properties of healthy WM, namely a constant 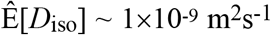 and a high anisotropy 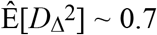. The anisotropy metric 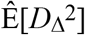 is found to be unaffected by the presence of fibre crossings (see lower right panel of **Figure 5**); this is in contrast to the widely popular Fractional Anisotropy (FA) metric, which is highly dependent on the degree of orientational order (Basser & Pierpaoli, 1996). Significantly lower 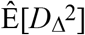 values are found at WM-GM interfaces, an observation that indicates the presence of tissue components with a lower diffusion anisotropy than that of components within pure WM voxels. Finally, we would like to note that glyphs close to ventricles do not reveal an increased *D*_iso_ or decreased *R*_2_, thus evidencing the successful resolution of signal contributions from distinct tissue components.

**Figure 5.**
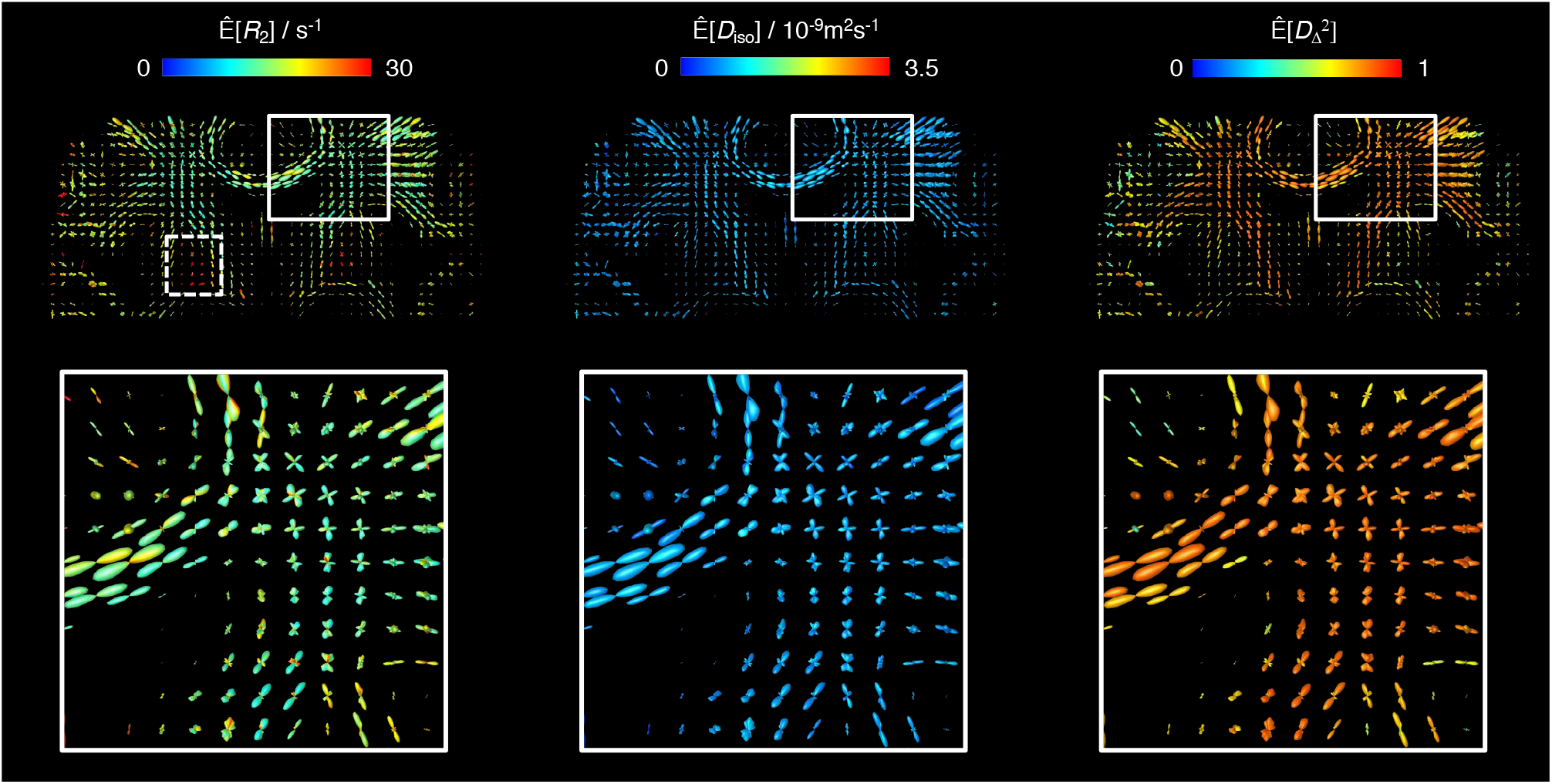
Orientation Distribution Function (ODF) maps coloured according to orientation-resolved means, 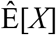, of *R*_2_, isotropic diffusivity *D*_iso_, and squared normalized diffusion anisotropy *D*_Δ_^2^. All 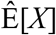 were calculated using Eq. (9) and are displayed on a linear scale. The lower panel displays a zoom into a region containing fibre crossings between the corpus callosum and the corticospinal tract. The dashed-line box in the top-left map identifies the high-*R*_2_ fibres found in the pallidum.

Focusing on the 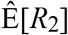-coloured ODFs shown in the left side panels of **Figure 5**, we find a population of fibres with considerably high 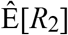 values in the midbrain region (see dashed box in the top left map of **Figure 5**). The fast-relaxing ODFs can be attributed to the myelinated axons that traverse the *globus pallidus*, an iron-rich basal ganglia structure that is characterized by particularly high *R*_2_ values (Hasan, Walimuni, Kramer, & Narayana, 2012; Knight et al., 2015). Not accounting for their significantly different *R*_2_ would then lead to an underestimation of the signal fraction of those high-*R*_2_ anisotropic components. Moreover, acquiring diffusion-weighted data measured at a single relatively high *τ*_E_ could even obscure the presence of anisotropic tissues in the *pallidum*.

To explore a possible angular dependence of the *R*_2_-**D** metrics from different fibre populations, we computed peak-specific 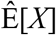 values and compared them against their respective *θ* coordinates. The results are displayed in **Figure 6**, where no clear relationship between 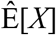 and peak orientation was observed for any of the extracted metrics. This observation is in contrast with previous *in vivo* brain MRI studies where a relationship between the *R*_2_ rates of myelinated fibres and their orientation relative to ***B***_0_ (Gil et al., 2016; Knight et al., 2015; McKinnon & Jensen, 2019) has been reported. In particular, Gil and co-workers (Gil et al., 2016) have estimated an angular variation of Δ*R*_2_ ~ 1.5 s^−1^ for healthy WM tissue. **Figure 3A** shows that the uncertainty of the Monte Carlo analysis procedure can introduce *R*_2_ differences of up to Δ*R*_2_ ~ 2.4 s^−1^ within a single fibre population; this suggests that a subtle *R*_2_ variation is very challenging to resolve with our minimally constrained approach and explains the approximately constant trend observed in **Figure 6** for the 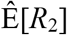 and 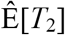 metrics. To better assess the minimum *R*_2_ uncertainty of our analysis protocol, we focused on individual CC voxels yielding single-lobe ODFs and computed the interquartile range of mean *R*_2_ values computed from different bootstrap 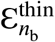 components. Such procedure yielded interquartile ranges of 1.5–2.5 s^−1^ for the various CC voxels, providing further evidence that the *R*_2_ uncertainty of a single fibre population determined through the Monte Carlo inversion is on the same order of magnitude as the *R*_2_ orientational effects reported in ref. (Gil et al., 2016).

**Figure 6.**
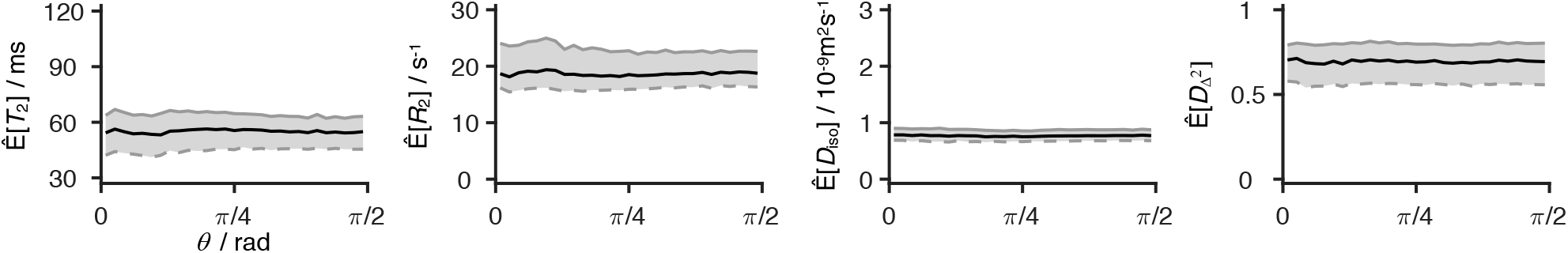
Peak-specific means, 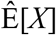, of *T*_2_, *R*_2_, isotropic diffusivity *D*_iso_, and squared normalized diffusion anisotropy *D*_Δ_^2^ plotted as a function of *θ*, the polar angle describing the orientation of the various peaks relative to the laboratory frame of reference. The peaks were estimated from local maxima of the smooth ODF, as described in the *Methods* section. The *θ* angles were sorted into 30 different bins. The solid grey, solid black, and dashed grey lines represent the 75^th^, 50^th^ (or median), and 25^th^ percentile, respectively, of each angle bin. The shaded grey region illustrates the interquartile range of each angle bin.

Despite the fact that no global *R*_2_(*θ*) behaviour could be teased out, the proposed method allowed the detection of relaxation differences between distinct WM tracts. As shown in **Figure 7**, these differences are best visualized in a *T*_2_ scale spanning a more constrained interval of values than the *R*_2_ scale used in **Figure 5**. Inspection of **Figure 7** reveals that both the CST and the *forceps major* tracts are characterized by considerably higher 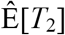 (lower 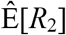) values. These observations are in accordance with the results of (Lampinen et al., 2020), where larger *T*_2_ times were consistently found in the CST. The higher *T*_2_ of the CST is also observed in voxels containing fibre crossings, with 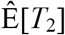 differences being discerned between the ODF lobes corresponding to the CST and the lobes that capture fibre populations from other tracts (see bottom right panels of **Figure 7**). While the exact mechanisms driving the high *T*_2_ values found in the CST and the forceps major are still unclear, it is worth mentioning that these tracts are known to feature higher-than-average fractions of both myelin water (Coelho, Pozo, Jespersen, & Frangi, 2019) and large axons (Dell’Acqua et al., 2019).

**Figure 7.**
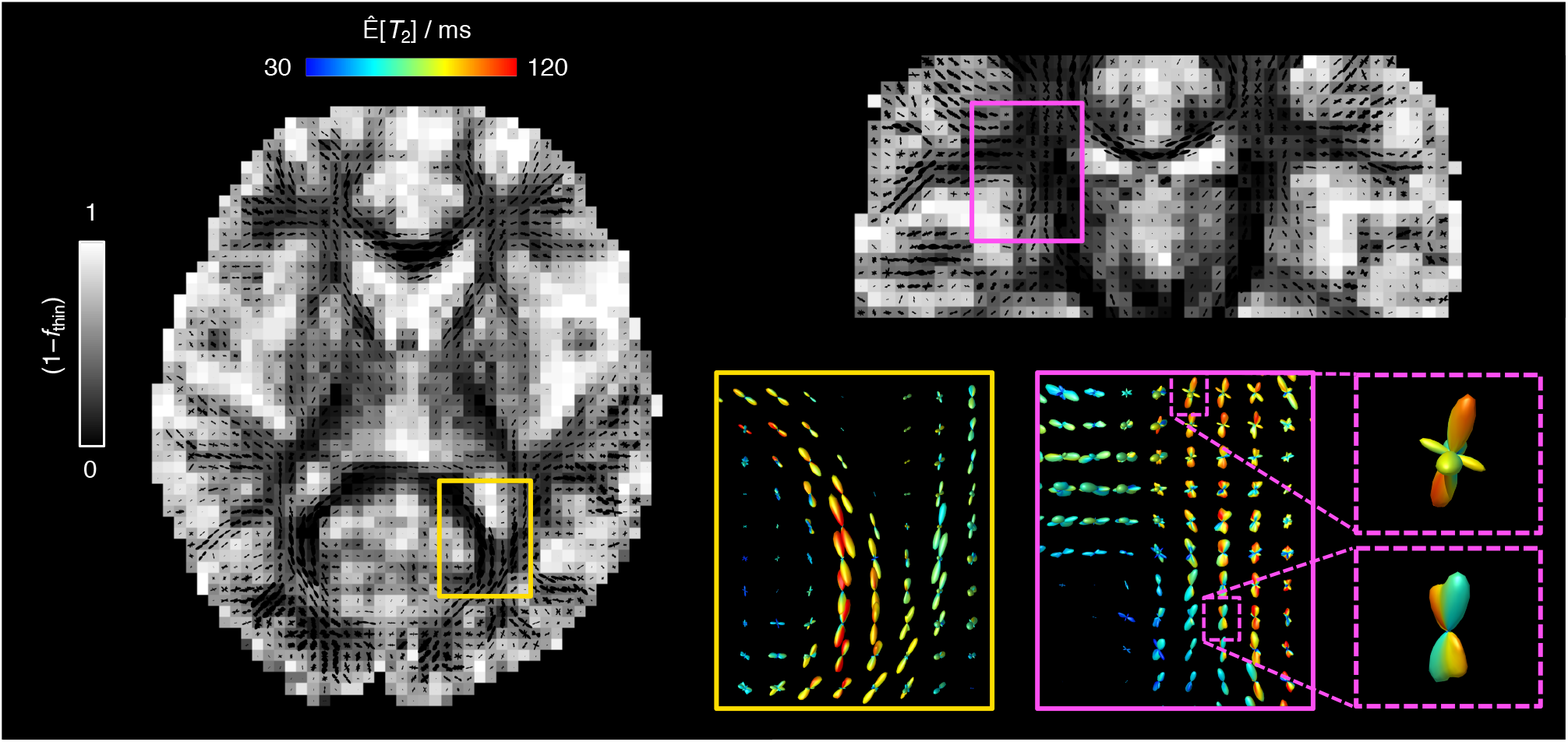
Orientation Distribution Function (ODF) maps coloured according to the orientation-resolved means of *T*_2_, 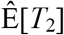. The 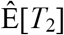 values are displayed on a linear colour scale. The left and topright panels display the sets of ODF glyphs superimposed on a grey-scaled map showing the signal fractions from the ‘big’ and ‘thick’ bin populations (1 – *f*_thin_) (non-fibre-like components). The zoom-ins in the lower-right panel offer a more detailed look into selected regions (continuous line boxes) and voxels (dashed line boxes) containing crossing between fibre populations with distinct 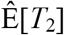. The observed high-*T*_2_ components are assigned to the forceps major (yellow boxes) and the corticospinal tract (magenta boxes).

While useful for visualization purposes, the colour-coded glyphs derived in this work are however impractical for quantifying the dispersion of *R*_2_-**D** descriptors within a given ODF lobe. For example, the *in silico* distributions from **Figure 3A** demonstrate that a single fibre population may comprise a dispersion of *T*_2_ values that cannot be accounted for by simply computing the 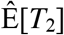 value of the associated ODF peak. In this regard, we suggest using the ODFs and corresponding peaks as a guide to define additional bins in the (*θ,ϕ*) space and to subsequently assign the voxel-wise 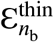 components into the various orientation-resolved bins. Once the orientation bins have been defined and the 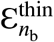 components assigned, orientation-specific statistical metrics and uncertainty measures can be estimated by exploring the variability of components within a given (*θ*,*ϕ*)-bin. An illustration of this procedure is presented in **Figure 8** for an SLF voxel comprising two crossing fibres. There, the (*θ*,*ϕ*)-space was divided into four quadrants centred around the extracted ODF peaks; average and dispersion measures were then calculated as the median and interquartile range of the 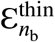 components falling within each quadrant.

**Figure 8.**
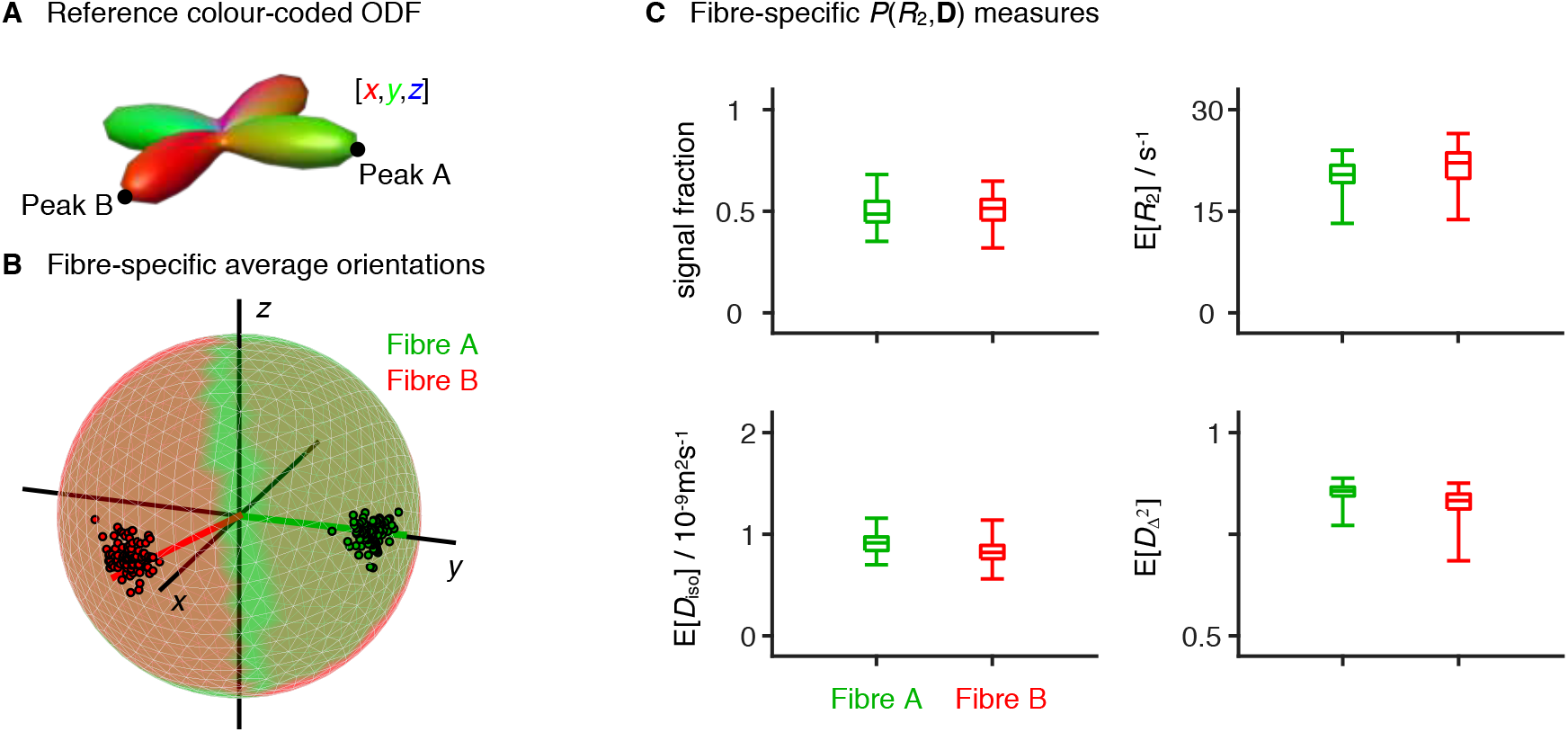
Orientation-resolved metrics estimated for a two-fibre-crossing voxel in the superior longitudinal fasciculus. (A) Orientation Distribution Function (ODF) estimated for the selected voxel. The black points identify the two peaks of the displayed ODF, peaks A and B. (B-C) Fibre-specific *R*_2_-**D** metrics. The (*θ*,*ϕ*) orientation space was divided into four quadrants centred on A, B, and their corresponding antipodes; ‘thin’ *R*_2_-**D** components, 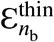, were then assigned to either fibre population A or fibre population B depending on their (*θ,ϕ*) coordinates (*e.g*. 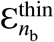 components falling into the quadrant centred on peak A, are assigned to fibre population A). For each orientation bin and each bootstrap, we estimate the mean signal fraction, *R*_2_, isotropic diffusivity *D*_iso_, squared normalized diffusion anisotropy *D*_Δ_^2^, and orientation, thus obtaining a set of 96×5 scalars: 96 different estimates of five distinct parameters. (B) Ensemble of fibre-resolved orientations displayed on the unit sphere. The colouring of the sphere identifies the (*θ*,*ϕ*) space assigned to each fibre population. The coloured lines indicate the peak orientation of fibres A (green) and B (red), while the black lines indicate the [*x*,*y*,*z*] coordinates. (C) Boxplots displaying the average and dispersion of the fibre-resolved signal fractions, *R*_2_, *D*_iso_, and *D*_Δ_^2^. The average was estimated as the median, while dispersion was assessed as the interquartile range. The whiskers identify the maximum and minimum estimated values.

The procedure depicted in **Figure 8** showcases the potential of using *P*(*R*_2_,**D**) distributions to extract the average and variance of fibre-specific metrics. In a preliminary work (Reymbaut, de Almeida Martins, et al., 2020), we combine the presented ODF framework with density-based clustering algorithms (Rodriguez & Laio, 2014) in order to sort 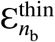 into different fibre populations and then calculate fibre-specific statistical metrics from the clustered *P*(*R*_2_,**D**) components.

## 4 CONCLUSION

This work presents analysis protocols to estimate and visualize orientation-resolved *R*_2_-**D** metrics in the living human brain. We build on a recently developed 5D relaxation-diffusion correlation framework where sub-voxel heterogeneity is resolved with nonparametric *P*(*R*_2_,**D**) distributions (de Almeida Martins et al., 2020), and convert the retrieved distributions to ODF glyphs informing on the relaxation-diffusion features along different orientations by mapping discrete *P*(*R*_2_,**D**) components to a dense mesh of (*θ*,*ϕ*) bins. Directionally-coloured ODFs estimated in such way were observed to capture fibre crossings in major WM tracts such as the CC, the CST, or the SLF. Similarly, arrays of *T*_2_-, *R*_2_-, *D*_iso_-, and *D*_Δ_^2^-coloured ODF glyphs were observed to facilitate a clean and compact visualization of the *R*_2_-**D** properties of anisotropic tissues. Maps of relaxation-coloured ODF also enabled the identification of fast-relaxing anisotropic components in the *globus pallidus* and the observation of long *T*_2_ times in the CST and the *forceps major*.

The proposed framework relies 5D *R*_2_-**D** distributions that provide a clean 3D mapping of the signal contributions from different sub-voxel tissue environments and allow the estimation of relaxation or diffusion differences between distinct fibre populations. Moreover, the *P*(*R*_2_,**D**) are retrieved from the data without the need to *a priori* fix signal response functions or formulating assumptions about the number of microscopic tissue components. This is in contrast with traditional (Anderson, 2005; Dell’Acqua et al., 2007; Dell’Acqua & Tournier, 2019; Jian & Vemuri, 2007; Tournier et al., 2007; Tournier et al., 2004) or multi-tissue (Jeurissen et al., 2014) spherical deconvolution approaches, which assume a single response function for WM tissue and do not accommodate for microstructural differences across fibres. The caveat is that the proposed method hinges on signal acquisition in a high dimensional space in order to better capture the signal contrast between environments with different MR properties (Topgaard, 2019); a comprehensive sampling of such space in turn introduces acquisition times that are longer than the ones currently used in spherical deconvolution protocols. However, there is potential to reduce the scan time either by using multi-band acquisition schemes (Barth, Breuer, Koopmans, Norris, & Poser, 2016) or designing more abbreviated acquisition protocols. Recent advances in nonparametric protocol optimization (Bates, Daducci, & Caruyer, 2019; Song & Xiao, 2020) are expected to facilitate a reduction of the measured data points while keeping a good performance of the Monte Carlo inversion procedure. Protocol optimization strategies can additionally be used to maximise the angular coverage of the acquisition scheme and hopefully increase the angular resolution of the retrieved distributions (Caruyer et al., 2011).

The information retrieved with the presented methodology can serve as an input for fibre tracking algorithms and used to extract individual WM pathways. If combined with tractometry frameworks (Bells et al., 2011; Chamberland et al., 2019; De Santis et al., 2014; Rheault et al., 2017; Yeatman et al., 2012), the correlations across the *R*_2_-**D** space would allow a comprehensive inspection of the relaxation and diffusion properties along a given WM tract. Since no universal signal response kernels are assumed, microstructural differences between tracts can be investigated and teased out. This feature is particularly promising for clinical research studies (Fornito, Zalesky, & Breakspear, 2015) where the 5D *R*_2_-**D** correlation framework could be used to investigate pathology induced changes along specific WM bundles.

## ACKNOWLEDGEMENTS

This work was financially supported by the Swedish Foundation for Strategic Research (AM13-0090 and ITM17-0267) and the Swedish Research Council (2014-3910 and 2018-03697). D. K. Jones and C. M. W. Tax were supported by a Wellcome Trust Investigator Award (096646/Z/11/Z), C. M. W. Tax by a Sir Henry Wellcome Fellowship and a Veni grant (17331) from the Dutch Research Council (NWO), and D. K. Jones by a Wellcome Trust Strategic Award (104943/Z/14/Z).

## REFERENCES

Aboitiz, F., Scheibel, A. B., Fisher, R. S., & Zaidel, E. (1992). Fiber composition of the human corpus callosum. Brain Research, 598(1-2), 143–153.

Alexander-Bloch, A., Giedd, J. N., & Bullmore, E. (2013). Imaging structural co-variance between human brain regions. Nature Reviews Neuroscience, 14(5), 322–336.

Anderson, A. W. (2005). Measurement of fiber orientation distributions using high angular resolution diffusion imaging. Magnetic Resonance in Medicine, 54(5), 1194–1206.

Andersson, J. L., Skare, S., & Ashburner, J. (2003). How to correct susceptibility distortions in spinecho echo-planar images: application to diffusion tensor imaging. Neuroimage, 20(2), 870–888.

Assaf, Y. (2019). Imaging laminar structures in the gray matter with diffusion MRI. NeuroImage, 197, 677–688.

Bak, M., & Nielsen, N. C. (1997). REPULSION, a novel approach to efficient powder averaging in solid-state NMR. Journal of Magnetic Resonance, 125(1), 132–139.

Barnea-Goraly, N., Kwon, H., Menon, V., Eliez, S., Lotspeich, L., & Reiss, A. L. (2004). White matter structure in autism: preliminary evidence from diffusion tensor imaging. Biological Psychiatry, 55(3), 323–326.

Barth, M., Breuer, F., Koopmans, P. J., Norris, D. G., & Poser, B. A. (2016). Simultaneous multislice (SMS) imaging techniques. Magnetic Resonance in Medicine, 75(1), 63–81.

Basser, P. J., Pajevic, S., Pierpaoli, C., Duda, J., & Aldroubi, A. (2000). In vivo fiber tractography using DT-MRI data. Magnetic Resonance in Medicine, 44(4), 625–632.

Basser, P. J., & Pierpaoli, C. (1996). Microstructural and physiological features of tissues elucidated by quantitative-diffusion-tensor MRI. Journal of magnetic resonance, Series B, 111(3), 209–219.

Bates, A., Daducci, A., & Caruyer, E. (2019). Multi-Dimensional Diffusion MRI Sampling Scheme: B-tensor Design and Accurate Signal Reconstruction. Paper presented at the 27th Annual Meeting of the ISMRM, Montreal, Canada.

Bells, S., Cercignani, M., Deoni, S., Assaf, Y., Pasternak, O., Evans, C., … Jones, D. K. (2011). Tractometry–comprehensive multi-modal quantitative assessment of white matter along specific tracts. Paper presented at the 19th Annual Meeting of the ISMRM, Montreal, Canada.

Benjamini, D., & Basser, P. J. (2016). Use of marginal distributions constrained optimization (MADCO) for accelerated 2D MRI relaxometry and diffusometry. Journal of Magnetic Resonance, 271, 40–45.

Benjamini, D., & Basser, P. J. (2017). Magnetic resonance microdynamic imaging reveals distinct tissue microenvironments. NeuroImage, 163, 183–196.

Berman, P., Levi, O., Parmet, Y., Saunders, M., & Wiesman, Z. (2013). Laplace inversion of low-resolution NMR relaxometry data using sparse representation methods. Concepts in Magnetic Resonance Part A, 42(3), 72–88.

Brodmann, K. (1909). Vergleichende Lokalisationslehre der Grosshirnrinde in ihren Prinzipien dargestellt auf Grund des Zellenbaues: Barth.

Callaghan, P. T., & Stepišnik, J. (1996). Generalized analysis of motion using magnetic field gradients. In Advances in magnetic and optical resonance (Vol. 19, pp. 325–388): Elsevier.

Caruyer, E., Cheng, J., Lenglet, C., Sapiro, G., Jiang, T., & Deriche, R. (2011, 2011-09-22). Optimal Design of Multiple Q-shells experiments for Diffusion MRI. Paper presented at the MICCAI Workshop on Computational Diffusion MRI -CDMRI’11, Toronto, Canada.

Chamberland, M., Raven, E. P., Genc, S., Duffy, K., Descoteaux, M., Parker, G. D., … Jones, D. K. (2019). Dimensionality reduction of diffusion MRI measures for improved tractometry of the human brain. NeuroImage, 200, 89–100.

Coelho, S., Pozo, J. M., Jespersen, S. N., & Frangi, A. F. (2019, 2019//). Optimal Experimental Design for Biophysical Modelling in Multidimensional Diffusion MRI. Paper presented at the Medical Image Computing and Computer Assisted Intervention – MICCAI 2019, Cham.

Daducci, A., Canales-Rodríguez, E. J., Zhang, H., Dyrby, T. B., Alexander, D. C., & Thiran, J.-P. (2015). Accelerated Microstructure Imaging via Convex Optimization (AMICO) from diffusion MRI data. NeuroImage, 105, 32–44.

de Almeida Martins, J. P., Tax, C. M. W., Szczepankiewicz, F., Jones, D. K., Westin, C.-F., & Topgaard, D. (2020). Transferring principles of solid-state and Laplace NMR to the field of in vivo brain MRI. Magn. Reson., 1(1), 27–43.

de Almeida Martins, J. P., & Topgaard, D. (2016). Two-Dimensional Correlation of Isotropic and Directional Diffusion Using NMR. Phys Rev Lett, 116(8), 087601.

de Almeida Martins, J. P., & Topgaard, D. (2018). Multidimensional correlation of nuclear relaxation rates and diffusion tensors for model-free investigations of heterogeneous anisotropic porous materials. Sci Rep, 8(1), 2488.

de Kort, D. W., van Duynhoven, J. P., Hoeben, F. J., Janssen, H. M., & Van As, H. (2014). NMR nanoparticle diffusometry in hydrogels: enhancing sensitivity and selectivity. Anal. Chem., 86(18), 9229–9235.

De Santis, S., Assaf, Y., Jeurissen, B., Jones, D. K., & Roebroeck, A. (2016). T1 relaxometry of crossing fibres in the human brain. NeuroImage, 141, 133–142.

De Santis, S., Drakesmith, M., Bells, S., Assaf, Y., & Jones, D. K. (2014). Why diffusion tensor MRI does well only some of the time: Variance and covariance of white matter tissue microstructure attributes in the living human brain. NeuroImage, 89, 35–44.

Dell’Acqua, F., Dallyn, R., Chiappiniello, A., Beyh, A., Tax, C. M. W., Jones, D. K., & Catani, M. (2019). Temporal Diffusion Ratio (TDR): A Diffusion MRI technique to map the fraction and spatial distribution of large axons in the living human brain. Paper presented at the 27th Annual Meeting of the ISMRM, Montreal, Canada.

Dell’Acqua, F., Rizzo, G., Scifo, P., Clarke, R. A., Scotti, G., & Fazio, F. (2007). A Model-Based Deconvolution Approach to Solve Fiber Crossing in Diffusion-Weighted MR Imaging. IEEE Trans. Biomed. Eng., 54, 462–472.

Dell’Acqua, F., & Tournier, J.-D. (2019). Modelling white matter with spherical deconvolution: How and why? NMR Biomed, 32(4), e3945.

Does, M. D. (2018). Inferring brain tissue composition and microstructure via MR relaxometry. NeuroImage, 182, 136–148.

Eriksson, S., Lasič, S., Nilsson, M., Westin, C.-F., & Topgaard, D. (2015). NMR diffusion-encoding with axial symmetry and variable anisotropy: Distinguishing between prolate and oblate microscopic diffusion tensors with unknown orientation distribution. J Chem Phys, 142(10), 104201.

Fornito, A., Zalesky, A., & Breakspear, M. (2015). The connectomics of brain disorders. Nature Reviews Neuroscience, 16(3), 159–172.

Gil, R., Khabipova, D., Zwiers, M., Hilbert, T., Kober, T., & Marques, J. P. (2016). An in vivo study of the orientation-dependent and independent components of transverse relaxation rates in white matter. NMR in Biomedicine, 29(12), 1780–1790.

Guo, F., Tax, C. M. W., Luca, A. D., Viergever, M. A., Heemskerk, A., & Leemans, A. (2019). Effects of Inaccurate Response Function Calibration on Characteristics of the Fiber Orientation Distribution in Diffusion MRI. bioRxiv, 760546.

Hasan, K. M., Walimuni, I. S., Kramer, L. A., & Narayana, P. A. (2012). Human brain iron mapping using atlas-based T2 relaxometry. Magnetic Resonance in Medicine, 67(3), 731–739.

Henkelman, R. M., Stanisz, G. J., Kim, J. K., & Bronskill, M. J. (1994). Anisotropy of NMR properties of tissues. Magnetic Resonance in Medicine, 32(5), 592–601.

Howard, A. F. D., Mollink, J., Kleinnijenhuis, M., Pallebage-Gamarallage, M., Bastiani, M., Cottaar, M., … Jbabdi, S. (2019). Joint modelling of diffusion MRI and microscopy. NeuroImage, 201, 116014.

Jelescu, I. O., Veraart, J., Fieremans, E., & Novikov, D. S. (2016). Degeneracy in model parameter estimation for multi-compartmental diffusion in neuronal tissue. NMR Biomed, 29(1), 33–47.

Jespersen, S. N., Lundell, H., Sønderby, C. K., & Dyrby, T. B. (2013). Orientationally invariant metrics of apparent compartment eccentricity from double pulsed field gradient diffusion experiments. NMR in Biomedicine, 26(12), 1647–1662.

Jeurissen, B., Tournier, J.-D., Dhollander, T., Connelly, A., & Sijbers, J. (2014). Multi-tissue constrained spherical deconvolution for improved analysis of multi-shell diffusion MRI data. NeuroImage, 103, 411–426.

Jian, B., & Vemuri, B. C. (2007). A Unified Computational Framework for Deconvolution to Reconstruct Multiple Fibers From Diffusion Weighted MRI. IEEE Transactions on Medical Imaging, 26(11), 1464–1471.

Jian, B., Vemuri, B. C., Özarslan, E., Carney, P. R., & Mareci, T. H. (2007). A novel tensor distribution model for the diffusion-weighted MR signal. NeuroImage, 37(1), 164–176.

Jones, D. K., Horsfield, M. A., & Simmons, A. (1999). Optimal strategies for measuring diffusion in anisotropic systems by magnetic resonance imaging. Magnetic Resonance in Medicine, 42(3), 515–525.

Kaden, E., Kelm, N. D., Carson, R. P., Does, M. D., & Alexander, D. C. (2016). Multi-compartment microscopic diffusion imaging. NeuroImage, 139, 346–359.

Kellner, E., Dhital, B., Kiselev, V. G., & Reisert, M. (2016). Gibbs-ringing artifact removal based on local subvoxel-shifts. Magnetic Resonance in Medicine, 76(5), 1574–1581.

Kindlmann, G. (2004). Superquadric tensor glyphs. Paper presented at the Proceedings of the Sixth Joint Eurographics-IEEE TCVG conference on Visualization.

Klein, S., Staring, M., Murphy, K., Viergever, M. A., & Pluim, J. P. (2009). Elastix: a toolbox for intensity-based medical image registration. IEEE Transactions on Medical Imaging, 29(1), 196–205.

Knight, M. J., Wood, B., Couthard, E., & Kauppinen, R. (2015). Anisotropy of spin-echo T2 relaxation by magnetic resonance imaging in the human brain in vivo. Biomedical Spectroscopy and Imaging, 4(3), 299–310.

Kroeker, R. M., & Mark Henkelman, R. (1986). Analysis of biological NMR relaxation data with continuous distributions of relaxation times. Journal of Magnetic Resonance (1969), 69(2), 218–235.

Lampinen, B., Szczepankiewicz, F., Mårtensson, J., van Westen, D., Hansson, O., Westin, C.-F., & Nilsson, M. (2020). Towards unconstrained compartment modeling in white matter using diffusion-relaxation MRI with tensor-valued diffusion encoding. Magnetic Resonance in Medicine, n/a(n/a).

Lasič, S., Szczepankiewicz, F., Eriksson, S., Nilsson, M., & Topgaard, D. (2014). Microanisotropy imaging: quantification of microscopic diffusion anisotropy and orientational order parameter by diffusion MRI with magic-angle spinning of the q-vector. Frontiers in Physics, 2, 11.

Lawrenz, M., Koch, M. A., & Finsterbusch, J. (2010). A tensor model and measures of microscopic anisotropy for double-wave-vector diffusion-weighting experiments with long mixing times. J Magn Reson, 202(1), 43–56.

Lawson, C. L., & Hanson, R. J. (1974). Solving least squares problems. Englewood Cliffs, NJ: Prentice-Hall.

Le Bihan, D. (1995). Molecular diffusion, tissue microdynamics and microstructure. NMR in Biomed., 8(7), 375–386.

Lebel, C., Walker, L., Leemans, A., Phillips, L., & Beaulieu, C. (2008). Microstructural maturation of the human brain from childhood to adulthood. NeuroImage, 40(3), 1044–1055.

Leemans, A. (2010). Visualization of diffusion MRI data. In D. K. Jones (Ed.), Diffusion MRI: Oxford University Press.

Lim, K. O., Hedehus, M., Moseley, M., de Crespigny, A., Sullivan, E. V., & Pfefferbaum, A. (1999). Compromised White Matter Tract Integrity in Schizophrenia Inferred From Diffusion Tensor Imaging. Archives of General Psychiatry, 56(4), 367–374.

Lindblom, G., Wennerström, H., & Arvidson, G. (1977). Translational diffusion in model membranes studied by nuclear magnetic resonance. International Journal of Quantum Chemistry, 12(2), 153_158.

Lundell, H., Nilsson, M., Dyrby, T. B., Parker, G. J. M., Cristinacce, P. L. H., Zhou, F.-L., … Lasič, S. (2019). Multidimensional diffusion MRI with spectrally modulated gradients reveals unprecedented microstructural detail. Scientific Reports, 9(1), 9026.

MacKay, A. L., Laule, C., Vavasour, I. M., Bjarnason, T., Kolind, S. H., & Mädler, B. (2006). Insights into brain microstructure from the T2 distribution. Magn Reson Imaging, 24(4), 515–525.

Mardia, K. V., & Jupp, P. E. (2009). Directional statistics (Vol. 494): John Wiley & Sons.

McKinnon, E. T., & Jensen, J. H. (2019). Measuring intra-axonal T2 in white matter with direction-averaged diffusion MRI. Magnetic Resonance in Medicine, 81(5), 2985–2994.

Mitchell, J., Chandrasekera, T. C., & Gladden, L. F. (2012). Numerical estimation of relaxation and diffusion distributions in two dimensions. Progress in Nuclear Magnetic Resonance Spectroscopy, 62, 34–50.

Mori, S., Crain, B. J., Chacko, V. P., & Van Zijl, P. C. (1999). Three-dimensional tracking of axonal projections in the brain by magnetic resonance imaging. Annals of Neurology, 45(2), 265–269.

Nilsson, M., Szczepankiewicz, F., Lampinen, B., Ahlgren, A., de Almeida Martins, J. P., Lasič, S., … Topgaard, D. (2018). An open-source framework for analysis of multidimensional diffusion MRI data implemented in MATLAB. Paper presented at the 26th Annual Meeting of the ISMRM, Paris, France.

Nilsson, M., Szczepankiewicz, F., van Westen, D., & Hansson, O. (2015). Extrapolation-Based References Improve Motion and Eddy-Current Correction of High B-Value DWI Data: Application in Parkinson’s Disease Dementia. PLOS ONE, 10(11), e0141825.

Ning, L., Gagoski, B., Szczepankiewicz, F., Westin, C. F., & Rathi, Y. (2020). Joint RElaxation-Diffusion Imaging Moments to Probe Neurite Microstructure. IEEE Transactions on Medical Imaging, 39(3), 668–677.

Novikov, D. S., Fieremans, E., Jespersen, S. N., & Kiselev, V. G. (2019). Quantifying brain microstructure with diffusion MRI: Theory and parameter estimation. NMR Biomed, 32(4), e3998.

Novikov, D. S., Veraart, J., Jelescu, I. O., & Fieremans, E. (2018). Rotationally-invariant mapping of scalar and orientational metrics of neuronal microstructure with diffusion MRI. NeuroImage, 174, 518–538.

Pajevic, S., & Pierpaoli, C. (2000). Color schemes to represent the orientation of anisotropic tissues from diffusion tensor data: application to white matter fiber tract mapping in the human brain. Magnetic Resonance in Medicine, 43(6), 921–921.

Parker, G. D., Marshall, D., Rosin, P. L., Drage, N., Richmond, S., & Jones, D. K. (2013). A pitfall in the reconstruction of fibre ODFs using spherical deconvolution of diffusion MRI data. NeuroImage, 65, 433–448.

Passingham, R. E., Stephan, K. E., & Kötter, R. (2002). The anatomical basis of functional localization in the cortex. Nature Reviews Neuroscience, 3(8), 606–616.

Peeters, T. H. J. M., Prckovska, V., Almsick, M. v., Vilanova, A., & Romeny, B. M. t. H. (2009, 20-23 April 2009). Fast and sleek glyph rendering for interactive HARDI data exploration. Paper presented at the 2009 IEEE Pacific Visualization Symposium.

Pierpaoli, C., Jezzard, P., Basser, P. J., Barnett, A., & Di Chiro, G. (1996). Diffusion tensor MR imaging of the human brain. Radiology, 201(3), 637–648.

Prange, M., & Song, Y.-Q. (2009). Quantifying uncertainty in NMR T2 spectra using Monte Carlo inversion. J Magn Reson, 196(1), 54–60.

Provencher, S. W. (1982). CONTIN: a general purpose constrained regularization program for inverting noisy linear algebraic and integral equations. Computer Physics Communications, 27(3), 229–242.

Reymbaut, A., de Almeida Martins, J. P., Tax, C. M. W., Szczepankiewicz, F., Jones, D. K., & Topgaard, D. (2020). Resolving orientation-specific diffusion-relaxation features via Monte-Carlo density-peak clustering in heterogeneous brain tissue. arXiv preprint arXiv:2004.08626.

Reymbaut, A., Mezzani, P., de Almeida Martins, J. P., & Topgaard, D. (2020). Accuracy and precision of statistical descriptors obtained from multidimensional diffusion signal inversion algorithms. NMR in Biomedicine, n/a(n/a), e4267.

Rheault, F., Houde, J.-C., & Descoteaux, M. (2017). Visualization, Interaction and Tractometry: Dealing with Millions of Streamlines from Diffusion MRI Tractography. Frontiers in Neuroinformatics, 11, 42.

Rodriguez, A., & Laio, A. (2014). Clustering by fast search and find of density peaks. Science, 344(6191), 1492–1496.

Scherrer, B., Schwartzman, A., Taquet, M., Sahin, M., Prabhu, S. P., & Warfield, S. K. (2016). Characterizing brain tissue by assessment of the distribution of anisotropic microstructural environments in diffusion-compartment imaging (DIAMOND). Magnetic Resonance in Medicine, 76(3), 963–977.

Schiavi, S., Pizzolato, M., Ocampo-Pineda, M., Canales-Rodriguez, E. J., Thiran, J.-P., & Daducci, A. (2019). Is it feasible to directly access the bundle’s specific myelin content, instead of averaging? A study with Microstructure Informed Tractography. Paper presented at the 27th Annual Meeting of the ISMRM, Montreal, Canada.

Schmidt-Rohr, K., & Spiess, H. W. (1994). Multidimensional solid-state NMR and polymers: Academic Press.

Schultz, T., & Kindlmann, G. (2010). A Maximum Enhancing Higher-Order Tensor Glyph. Computer Graphics Forum, 29(3), 1143–1152.

Schultz, T., & Vilanova, A. (2019). Diffusion MRI visualization. NMR in Biomedicine, 32(4), e3902.

Sjölund, J., Szczepankiewicz, F., Nilsson, M., Topgaard, D., Westin, C.-F., & Knutsson, H. (2015). Constrained optimization of gradient waveforms for generalized diffusion encoding. Journal of Magnetic Resonance, 261, 157–168.

Slator, P. J., Hutter, J., Palombo, M., Jackson, L. H., Ho, A., Panagiotaki, E., … Alexander, D. C. (2019). Combined diffusion-relaxometry MRI to identify dysfunction in the human placenta. Magn Reson Med, 82(1), 95–106.

Smith, S. M., Jenkinson, M., Woolrich, M. W., Beckmann, C. F., Behrens, T. E. J., Johansen-Berg, H., … Matthews, P. M. (2004). Advances in functional and structural MR image analysis and implementation as FSL. NeuroImage, 23, S208–S219.

Song, Y.-Q., & Xiao, L. (2020). Optimization of multidimensional MR data acquisition for relaxation and diffusion. NMR in Biomedicine, n/a(n/a), e4238.

Sporns, O., Tononi, G., & Kötter, R. (2005). The Human Connectome: A Structural Description of the Human Brain. PLOS Computational Biology, 1(4), e42.

Stejskal, E. O., & Tanner, J. E. (1965). Spin diffusion measurements: Spin echoes in the presence of a time-dependent field gradient. J. Chem. Phys., 42(1), 288–292.

Stepišnik, J. (1993). Time-dependent self-diffusion by NMR spin-echo. Physica B: Condensed Matter, 183(4), 343–350.

Szczepankiewicz, F., Lasič, S., Nilsson, M., Lundell, H., Westin, C.-F., & Topgaard, D. (2019). Is spherical diffusion encoding rotation invariant? An investigation of diffusion timedependence in the healthy brain. Paper presented at the 27th Annual Meeting of the ISMRM, Montreal, Canada.

Szczepankiewicz, F., Sjolund, J., Stahlberg, F., Latt, J., & Nilsson, M. (2019). Tensor-valued diffusion encoding for diffusional variance decomposition (DIVIDE): Technical feasibility in clinical MRI systems. PLOS ONE, 14(3), e0214238.

Szczepankiewicz, F., Westin, C.-F., & Nilsson, M. (2019). Maxwell-compensated design of asymmetric gradient waveforms for tensor-valued diffusion encoding. Magnetic Resonance Medicine, 82(4), 1424–1437.

Tax, C. M. W., Jeurissen, B., Vos, S. B., Viergever, M. A., & Leemans, A. (2014). Recursive calibration of the fiber response function for spherical deconvolution of diffusion MRI data. NeuroImage, 86, 67–80.

Topgaard, D. (2017). Multidimensional diffusion MRI. J Magn Reson, 275, 98–113.

Topgaard, D. (2019). Diffusion tensor distribution imaging. NMR Biomed, 32(5), e4066.

Tournier, J.-D. (2019). Diffusion MRI in the brain -Theory and concepts. Prog Nucl Magn Reson Spectrosc, 112-113, 1–16.

Tournier, J.-D., Calamante, F., & Connelly, A. (2007). Robust determination of the fibre orientation distribution in diffusion MRI: Non-negativity constrained super-resolved spherical deconvolution. NeuroImage, 35(4), 1459–1472.

Tournier, J.-D., Calamante, F., Gadian, D. G., & Connelly, A. (2004). Direct estimation of the fiber orientation density function from diffusion-weighted MRI data using spherical deconvolution. NeuroImage, 23(3), 1176–1185.

Tuch, D. S., Reese, T. G., Wiegell, M. R., Makris, N., Belliveau, J. W., & Wedeen, V. J. (2002). High angular resolution diffusion imaging reveals intravoxel white matter fiber heterogeneity. Magnetic Resonance Medicine, 48, 577–582.

Urbańczyk, M., Bernin, D., Koźmiński, W., & Kazimierczuk, K. (2013). Iterative thresholding algorithm for multiexponential decay applied to PGSE NMR data. Analytical chemistry, 85(3), 1828–1833.

Venkataramanan, L., Song, Y.-Q., & Hurlimann, M. D. (2002). Solving Fredholm integrals of the first kind with tensor product structure in 2 and 2.5 dimensions. IEEE Transactions on Signal Processing, 50(5), 1017–1026.

Veraart, J., Novikov, D. S., Christiaens, D., Ades-Aron, B., Sijbers, J., & Fieremans, E. (2016). Denoising of diffusion MRI using random matrix theory. NeuroImage, 142, 394–406.

Veraart, J., Novikov, D. S., & Fieremans, E. (2018). TE dependent Diffusion Imaging (TEdDI) distinguishes between compartmental T2 relaxation times. NeuroImage, 182, 360–369.

Vos, S. B., Tax, C. M., Luijten, P. R., Ourselin, S., Leemans, A., & Froeling, M. (2017). The importance of correcting for signal drift in diffusion MRI. Magnetic Resonance in Medicine, 77(1), 285–299.

Watson, G. (1965). Equatorial distributions on a sphere. Biometrika, 52(1/2), 193–201.

Werring, D. J., Clark, C. A., Barker, G. J., Thompson, A. J., & Miller, D. H. (1999). Diffusion tensor imaging of lesions and normal-appearing white matter in multiple sclerosis. Neurology, 52(8), 1626.

Westin, C.-F., Knutsson, H., Pasternak, O., Szczepankiewicz, F., Özarslan, E., van Westen, D., … Nilsson, M. (2016). Q-space trajectory imaging for multidimensional diffusion MRI of the human brain. NeuroImage, 135, 345–362.

Westin, C.-F., Maier, S. E., Khidhir, B., Everett, P., Jolesz, F. A., & Kikinis, R. (1999, 1999//). Image Processing for Diffusion Tensor Magnetic Resonance Imaging. Paper presented at the Medical Image Computing and Computer Assisted Intervention – MICCAI 1999, Berlin, Heidelberg.

Whittall, K. P., & MacKay, A. L. (1989). Quantitative interpretation of NMR relaxation data. Journal of Magnetic Resonance, 84(1), 134–152.

Whittall, K. P., MacKay, A. L., Graeb, D. A., Nugent, R. A., Li, D. K. B., & Paty, D. W. (1997). In vivo measurement of T2 distributions and water contents in normal human brain. Magnetic Resonance in Medicine, 37(1), 34–43.

Yeatman, J. D., Dougherty, R. F., Myall, N. J., Wandell, B. A., & Feldman, H. M. (2012). Tract Profiles of White Matter Properties: Automating Fiber-Tract Quantification. PLOS ONE, 7(11), e49790.

Zilles, K., & Amunts, K. (2010). Centenary of Brodmann’s map--conception and fate. Nat Rev Neurosci, 11(2), 139–145.

